# Separation of transcriptional repressor and activator functions in HDAC3

**DOI:** 10.1101/2022.06.11.495646

**Authors:** Min Tang, Isabel Regadas, Sergey Belikov, Olga Shilkova, Lei Xu, Erik Wernersson, Xuewen Liu, Hongmei Wu, Magda Bienko, Mattias Mannervik

## Abstract

The histone deacetylase HDAC3 is associated with the NCoR/SMRT co-repressor complex and its canonical function is in transcriptional repression, but it can also activate transcription. Here we show that the repressor and activator functions of HDAC3 can be genetically separated in *Drosophila*. A lysine substitution in the N-terminus (K26A) disrupts its catalytic activity and activator function, whereas a combination of substitutions (HEBI) abrogating the interaction with SMRTER enhance repressor activity beyond wild-type in the early embryo. We conclude that the critical functions of HDAC3 in embryo development involve catalytic-dependent gene activation and non-enzymatic repression by several mechanisms, including tethering of loci to the nuclear periphery.

## Introduction

Precise regulation of gene expression is essential for cell specification and embryo development. Gene regulation largely relies on transcription factors that recruit co-regulators to facilitate association of RNA polymerase II with promoters, in part by modulating the chromatin structure (Cramer, 2019). Many co-regulators are associated with enzymatic activities that can modify histones and non-histone substrates, and protein acetylation and deacetylation in particular are intimately linked to gene regulation (Verdin and Ott, 2015). The histone deacetylase (HDAC) family of enzymes can be divided into four classes based on their catalytic mechanism and sequence homology (Seto and Yoshida, 2014). The class I HDACs include HDAC1, HDAC2, HDAC3 and HDAC8. HDAC3 associates with nuclear receptor co-repressor 1 (NCoR1 or NCoR) and silencing mediator of retinoic acid and thyroid hormone receptor (SMRT; also known as NCoR2), which are both nuclear receptor co-repressors (Lonard and O’Malley B, 2007). TBL1X, TBL1XR1 and GPS2 are also subunits of these complexes. HDAC3 is unique among class I HDACs, since its catalytic function requires physical interaction with a conserved domain in NCoR and SMRT, the deacetylase-activating domain (DAD) (Codina et al., 2005; Guenther et al., 2001). The HDAC3-DAD interaction and HDAC activity is regulated by inositol tetraphosphate (Millard et al., 2013; Watson et al., 2012).

HDAC3 is involved in a diverse set of developmental, physiological and metabolic processes through different molecular mechanism (Emmett and Lazar, 2019). In the mouse liver, HDAC3 represses metabolic and circadian genes through histone deacetylation (Kim et al., 2018; Knutson et al., 2008), whereas in oligodendrocyte precursors the transcription factor STAT3 is inactivated by deacetylation (Zhang et al., 2016). By contrast, in brown adipose tissue HDAC3 activates expression of the uncoupling protein 1 gene by deacetylating the co-activator protein peroxisome proliferator-activated receptor gamma co-activator 1 alpha (PGC1α) (Emmett et al., 2017). It can also activate neural genes in forebrain neurons by deacetylating the transcription factor FOXO3 (Nott et al., 2016).

Although global deletion of HDAC3 in mice is embryonic lethal owing to gastrulation defects (Bhaskara et al., 2008), mice with mutations in the DAD of both NCoR and SMRT are born at the expected Mendelian ratios, despite the lack of detectable HDAC3 enzymatic activity (You et al., 2013). Furthermore, HDAC3 represses cardiomyocyte differentiation by non-enzymatic mechanisms. It silences cardiac linage genes through targeting to the nuclear lamina and formation of lamina-associated domains associated with heterochromatin and H3K9me2 (Poleshko et al., 2017). In addition, the non-enzymatic activity of HDAC3 represses transforming growth factor-β1 (TGFβ1) signaling by recruiting the H3K27 methyltransferase Polycomb repressive complex 2 (PRC2) to prevent the development of cardiac fibrosis (Lewandowski et al., 2015). HDAC3 is also necessary for recruitment of PRC2 and silencing of the X-chromosome for dosage compensation in females (McHugh et al., 2015), and for male fertility and stratification of the embryonic epidermis independent of its enzymatic activity (Szigety et al., 2020; Yin et al., 2021). These observations suggest that HDAC3 also has important non-enzymatic functions.

In *Drosophila*, HDAC3 controls imaginal disc and body size through H4K16 deacetylation (Lv et al., 2012; Zhu et al., 2008), and contributes to repressor activity of the Snail transcription factor in the early embryo (Qi et al., 2008). HDAC3 also cooperates with the SWI/SNF chromatin remodeling Brahma complex to prevent dedifferentiation of larval brain intermediate neural progenitors into neuroblasts (Koe et al., 2014). In larval salivary glands, HDAC3 is a positive regulator of the *hsp70* gene (Achary et al., 2014). However, the molecular functions of HDAC3 in embryo patterning remain unknown.

In order to dissect theses functions, we generated HDAC3 point mutants modelled on mammalian HDAC3. Tyrosine 298 in mammalian HDAC3 is located within the active site and required for catalytic activity, whereas K25 is important for interaction with the DAD-domain (Lombardi et al., 2011; Sun et al., 2013; Watson et al., 2012). A K25A mutation disrupts the DAD interaction and reduces, but does not completely eliminate, binding to full length NCoR (Sun et al., 2013; You et al., 2013). Since other HDACs do not interact with NCoR/SMRT, HDAC3 was mutated in four potential interaction clusters that are divergent between HDAC3 and HDAC1, and named ‘‘HEBI’’ for HDAC3 with Enzyme and Binding activities Inactivated, because the HEBI mutant abolished the interaction with NCoR/SMRT (Sun et al., 2013). In this work, we made corresponding mutants in endogenous and transgenic *Drosophila* HDAC3 and investigated their effect on survival and embryo patterning. Interestingly, we found that the activator and repressor functions of HDAC3 could be separated by these mutations.

## Results

### HDAC3 catalytic activity is required for *Drosophila* viability

To investigate the functions of *Drosophila* HDAC3 in development, we generated three mutants that are expected to reduce its catalytic activity (Fig. 1A, S1A-C). The Y303F, K26A and HEBI mutants are modeled on mammalian HDAC3, where the corresponding Y298F, K25A and HEBI substitutions were shown to interfere with catalysis, reduce the interaction with the SMRT/NCoR DAD domain, and eliminate the interaction with full length SMRT/NCoR, respectively (Sun et al., 2013). We cloned HA-tagged wild-type and mutant *Drosophila* HDAC3 genomic regions containing the endogenous promoter, and transfected these constructs into S2 cells. After immunoprecipitation and elution, an HDAC assay was performed and deacetylase activity calculated after normalization of protein levels by Western blot (Fig. 1B and Fig S1D). The Y303F, K26A and HEBI mutants showed a comparable reduction in catalytic activity compared to wild-type HDAC3. Next, we investigated if the HDAC3 mutants could interact with SMRTER. We co-transfected HA-tagged HDAC3 and either a V5-tagged SMRTER DAD domain or V5-tagged full length SMRTER (Fig. 1C, D). As expected, the HEBI mutant strongly reduced the interaction with full-length SMRTER and with the DAD domain, whereas binding of the Y303F mutant to SMRTER was similar to wild-type. To our surprise, the K26A mutation did not disrupt the interaction with the DAD domain (Fig. 1C), but it did reduce binding to full-length SMRTER (Fig. 1D). This shows that reduced catalytic activity of HEBI is likely due to decreased interaction with SMRTER, whereas the K26A mutation may additionally disrupt catalytic activity by some other mechanism. The Western blot also shows that the point mutations do not affect expression or stability of HDAC3.

**Figure 1:**
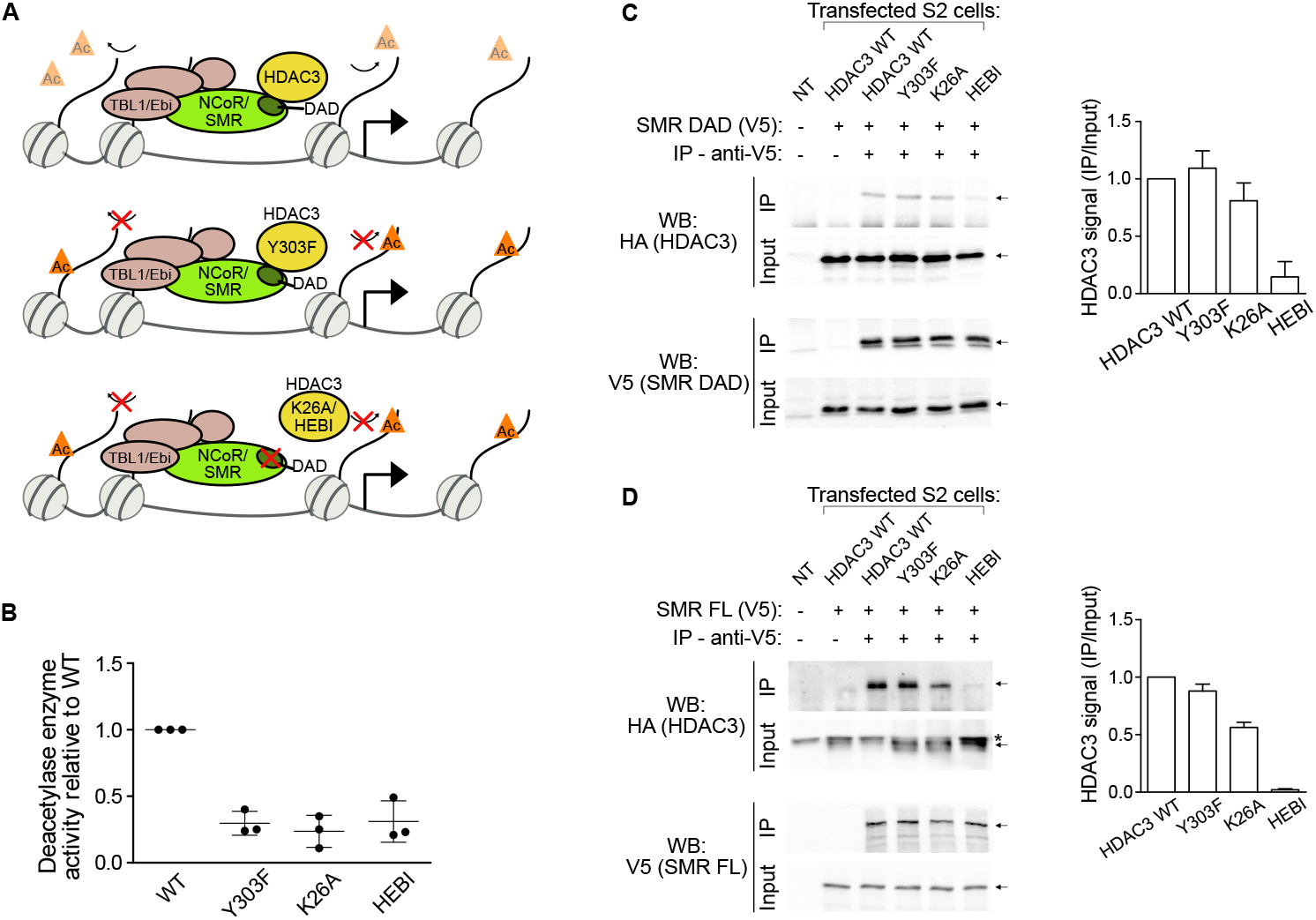
Impaired catalytic activity of mutant HDAC3 proteins. **A)** Scheme depicting wild-type and mutant (Y303F, K26A and HEBI) *Drosophila* HDAC3, their deacetylase capacity and interaction with NCoR/*Drosophila* SMRTER (SMR). **B)** Quantification of HDAC3 deacetylase activity after immunoprecipitation and normalization of protein levels by Western blot. **C)** and **D)** Western blots following co-immunoprecipitation experiments in S2 cells transfected with V5-tagged SMRTER DAD domain (**C**) or full length SMRTER (**D**) and HA-tagged wild-type or mutant HDAC3. The graphs show the HA-HDAC3 levels in the co-IP over the input (n=2-3, whiskers denote SD).

In order to explore catalytic and other possible functions for HDAC3 *in vivo*, we used homologous recombination to replace the HDAC3 transcription unit with a mini-*white* gene flanked by attP sites (Fig. S2A). The resulting flies, ΔHDAC3, are homozygous lethal and die during the pupal stage. However, 59% survive to the third instar larval stage (Table 1). It appears that the maternal contribution of wild-type HDAC3 allows for survival of homozygous mutant embryos and larvae to this stage. We then used a recombination-mediated cassette exchange (RMCE)-based approach to re-introduce either wild-type or the Y303F, K26A and HEBI mutants into the endogenous locus (Fig. S2A-C). Whereas the wild-type replaced flies (HDAC^WT^) develop into viable adults and 95% make it to third instar larvae, all three catalytically deficient mutants are homozygous lethal. Although many ΔHDAC3 and HEBI mutant animals reached the third instar stage, much fewer Y303F or K26A homozygous animals did (Table 1). This indicates that second site mutations on the Y303F and K26A chromosomes were inadvertently introduced during RMCE. Consistent with this notion, a larger fraction of trans-heterozygous combinations of ΔHDAC3 with Y303F or K26A survived to the third instar stage than the homozygous point mutants (Table 1), but they did not survive to adulthood. Western blots from three-day old larvae showed a 50% reduction of HDAC3 protein in heterozygous flies (Δ/HDAC^WT^) compared to a wild-type (*w*^*1118*^) strain (Fig. S2D). Similar levels were found in K26A trans-heterozygous mutants, whereas the Y303F and HEBI mutants had reduced HDAC3 protein levels. Some HDAC3 protein, presumably from the maternal load, remained in ΔHDAC3 homozygous larvae (Fig. S2D). This shows that the poor survival of K26A mutants cannot be explained by reduced protein expression, and therefore that the catalytic activity of HDAC3 is essential for viability.

**Table 1.**
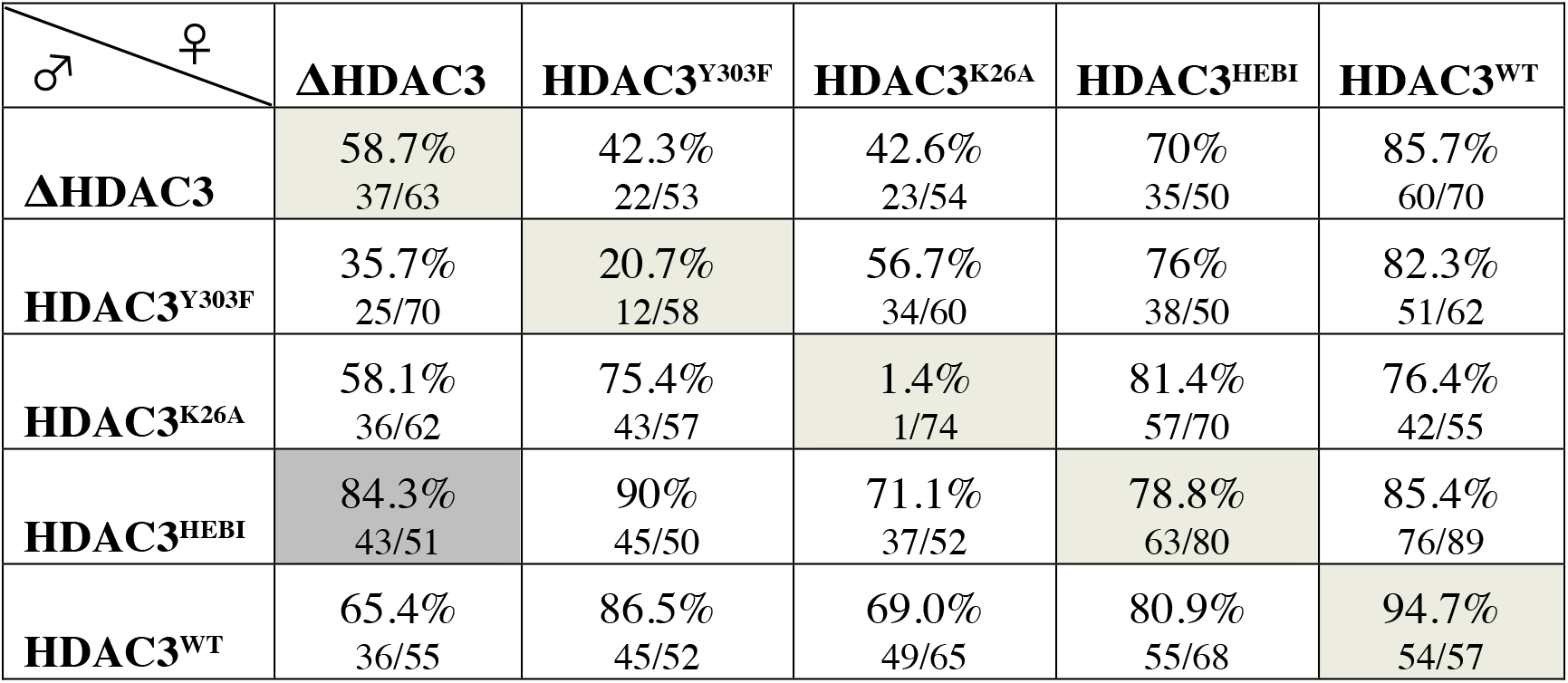
Survival to 3rd instar larvae

| ♂ \ ♀ | **ΔHDAC3** | **HDAC3<sup>Y303F</sup>** | **HDAC3<sup>K26A</sup>** | **HDAC3<sup>HEBI</sup>** | **HDAC3<sup>WT</sup>** |
| --- | --- | --- | --- | --- | --- |
| **ΔHDAC3** | 58.7%<br>37/63 | 42.3%<br>22/53 | 42.6%<br>23/54 | 70%<br>35/50 | 85.7%<br>60/70 |
| **HDAC3<sup>Y303F</sup>** | 35.7%<br>25/70 | 20.7%<br>12/58 | 56.7%<br>34/60 | 76%<br>38/50 | 82.3%<br>51/62 |
| **HDAC3<sup>K26A</sup>** | 58.1%<br>36/62 | 75.4%<br>43/57 | 1.4%<br>1/74 | 81.4%<br>57/70 | 76.4%<br>42/55 |
| **HDAC3<sup>HEBI</sup>** | 84.3%<br>43/51 | 90%<br>45/50 | 71.1%<br>37/52 | 78.8%<br>63/80 | 85.4%<br>76/89 |
| **HDAC3<sup>WT</sup>** | 65.4%<br>36/55 | 86.5%<br>45/52 | 69.0%<br>49/65 | 80.9%<br>55/68 | 94.7%<br>54/57 |

Animals from mothers with wild-type HDAC3 survived better than those from mutant mothers (Table 1, compare column 2 with column 6), showing that the maternal contribution influences their survival. The nature of the zygotic contribution is also important as animals from ΔHDAC3 heterozygous mothers survive better with a wt-replaced chromosome (65%) than with Y303F (36%) or K26A (58%, Table 1). Surprisingly, those with a paternal HEBI contribution survived better (84%) than those with a wt chromosome (Table 1), despite poor expression of the HEBI mutant protein (Fig. S2D). Thus, zygotic HEBI expression rescues viability better than wild-type HDAC3.

### HDAC3 catalytic function is dispensable for embryonic development

To understand the function of HDAC3 in embryo development, we used short hairpin RNA (shRNA)-mediated maternal knock-down of HDAC3, since the maternal contribution in ΔHDAC3 homozygous animals supports development to the larval/pupal stage. We generated shRNA-resistant rescue transgenes under control of the endogenous promoter and integrated them all at the same genomic landing site (Fig. 2A). As can be seen in Fig. 2B, maternal shRNA-mediated HDAC3 knock-down using the α-tubulin-Gal4VP16 driver results in a failure of most embryos to hatch into larvae. Introduction of one copy of a wild-type, shRNA-resistant transgene partially rescued this phenotype (Fig. 2B). A Western blot analysis showed an almost complete absence of HDAC3 protein in shRNA-depleted embryos, and incomplete restoration of HDAC3 protein expression in rescued embryos, which may explain the partial rescue (Fig. 2C). We then introduced the point mutations that disrupt HDAC3 catalytic activity in the shRNA-resistant rescue transgenes. The Y303F mutant rescued the hatching phenotype to the same extent as the wild-type transgene, whereas the K26A mutant failed to rescue (Fig. 2B). Surprisingly, the HEBI mutant rescued even better than the wild-type transgene (Fig. 2B). The Western blot shows that the K26A and HEBI proteins are expressed at levels similar to that from the wild-type transgene, whereas Y303F is more abundant (Fig. 2C). We performed RNA-seq from these embryos, which allowed us to distinguish between RNA expressed from the endogenous HDAC3 locus, and RNA from the shRNA-resistant transgenes. Consistent with the efficient depletion of HDAC3 protein, there is very little HDAC3 RNA in shRNA-treated embryos (Fig. 2D). In embryos from mothers that also have the wild-type shRNA-resistant transgene, HDAC3 levels are only partially restored and most of this RNA is derived from the transgene. Unexpectedly, in Y303F embryos there is both RNA derived from the shRNA-resistant transgene as well as plenty of RNA derived from the endogenous HDAC3 locus (Fig. 2D). This indicates that the Y303F protein feedbacks on the shRNA-mediated knockdown of HDAC3, and explains why the HDAC3 protein level is higher in these embryos than in wild-type rescued embryos (Fig. 2C). It also suggests that the reason why Y303F rescues the hatching phenotype to the same extent as the wild-type transgene is because wild-type HDAC3 protein is produced in these Y303F-rescued embryos (Fig. 2B). By contrast, in K26A embryos only mutant RNA is produced and the protein levels equal those in wild-type rescued embryos, but these embryos fail to hatch (Fig. 2B-D). This shows that the K26 residue is essential for embryo development. However, the RNA-seq cannot explain why the HEBI mutant rescues better than wild-type, as only shRNA-resistant RNA is detected (Fig. 2D). Thus, despite a loss in SMRTER interaction and catalytic activity (Fig. 1), the HEBI mutant supports embryonic development. Taken together, these results show that HDAC3 is required for embryo development, that catalytic activity is dispensable (since the HEBI mutant rescues the hatching phenotype), but that the K26A mutation disrupts the essential HDAC3 embryonic function.

**Figure 2:**
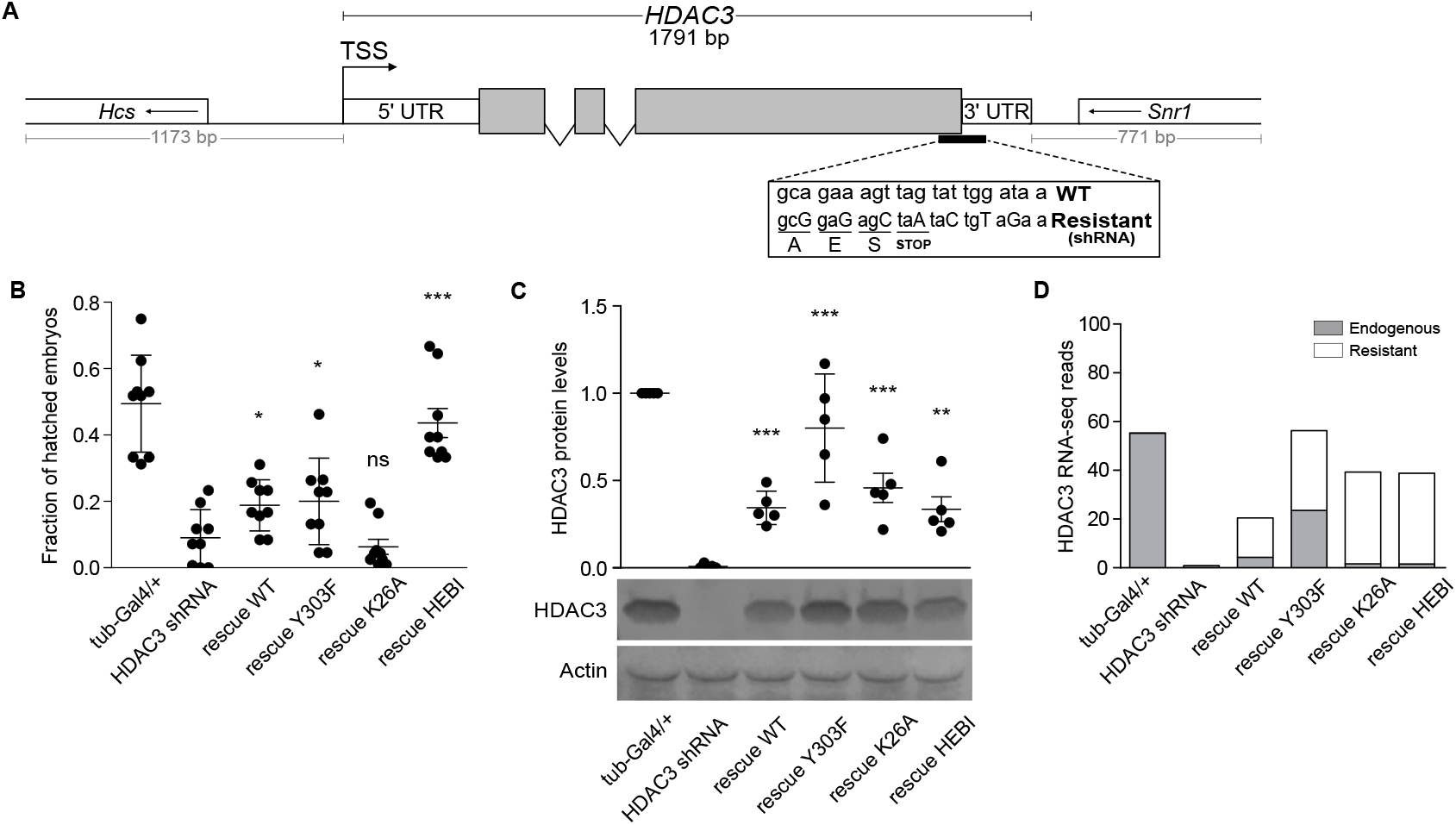
HDAC3^K26A^ disrupts but HDAC3^HEBI^ supports embryonic development. **A)** Schematic drawing of a HDAC3 shRNA-resistant rescue transgene under the control of its endogenous promoter. Wild-type and shRNA-resistant sequences are shown in the rectangle. **B)** Graph showing the fraction of control (tub-Gal4/+), maternal shRNA-mediated HDAC3 knocked-down (HDAC3 shRNA) and shRNA-resistant (rescue WT, rescue Y303F, rescue K26A and rescue HEBI) embryos that hatch into 1st instar larvae. **C)** Western blot showing the levels of HDAC3 protein normalized to Actin in HDAC3 shRNA and shRNA-resistant rescue embryos relative to Control (tub-GAL4/+). **D)** Endogenous and shRNA-resistant HDAC3 reads from RNA-seq in Control (tub-GAL4/+), HDAC3 shRNA and shRNA-resistant rescue embryos. Two-tailed, unpaired Student’s t test was applied to compare HDAC3 shRNA with shRNA-resistant animals. * - P<0.05, *** - P<0.0001. Error bars indicate SD.

### Unique functions for HDAC3 amino acids in embryonic patterning

We have previously demonstrated that specification of mesoderm in the early embryo by the Snail repressor requires the Ebi protein (Qi et al., 2008), a homolog of mammalian TBL1X/TBL1XR1. Although Ebi is part of the SMRTER complex and binds to HDAC3, the catalytic function of HDAC3 in Snail-mediated repression has not been investigated. We therefore collected 2-4 h old embryos and performed whole-mount in situ hybridization with a *short-gastrulation (sog)* probe (Fig. 3A). Snail protein is present in the mesoderm where it represses *sog* and restricts its expression to the presumptive neuroectoderm (Qi et al., 2008). In HDAC3 shRNA knock-down embryos, *sog* expression was partially de-repressed in the mesoderm, demonstrating that HDAC3 contributes to Snail-mediated repression of *sog* (Fig. 3A, B). The *sog* expression pattern was classified as normal, weak, or strong de-repression in the mesoderm (Fig. 3B). In wild-type rescued embryos, the amount of embryos with a normal *sog* pattern increased from 40% to 54%. Remarkably, all three HDAC3 mutant constructs also rescued *sog* expression in the mesoderm (Fig. 3B). This shows that HDAC3 catalytic activity is dispensable for Snail-mediated repression of *sog* transcription.

**Figure 3:**
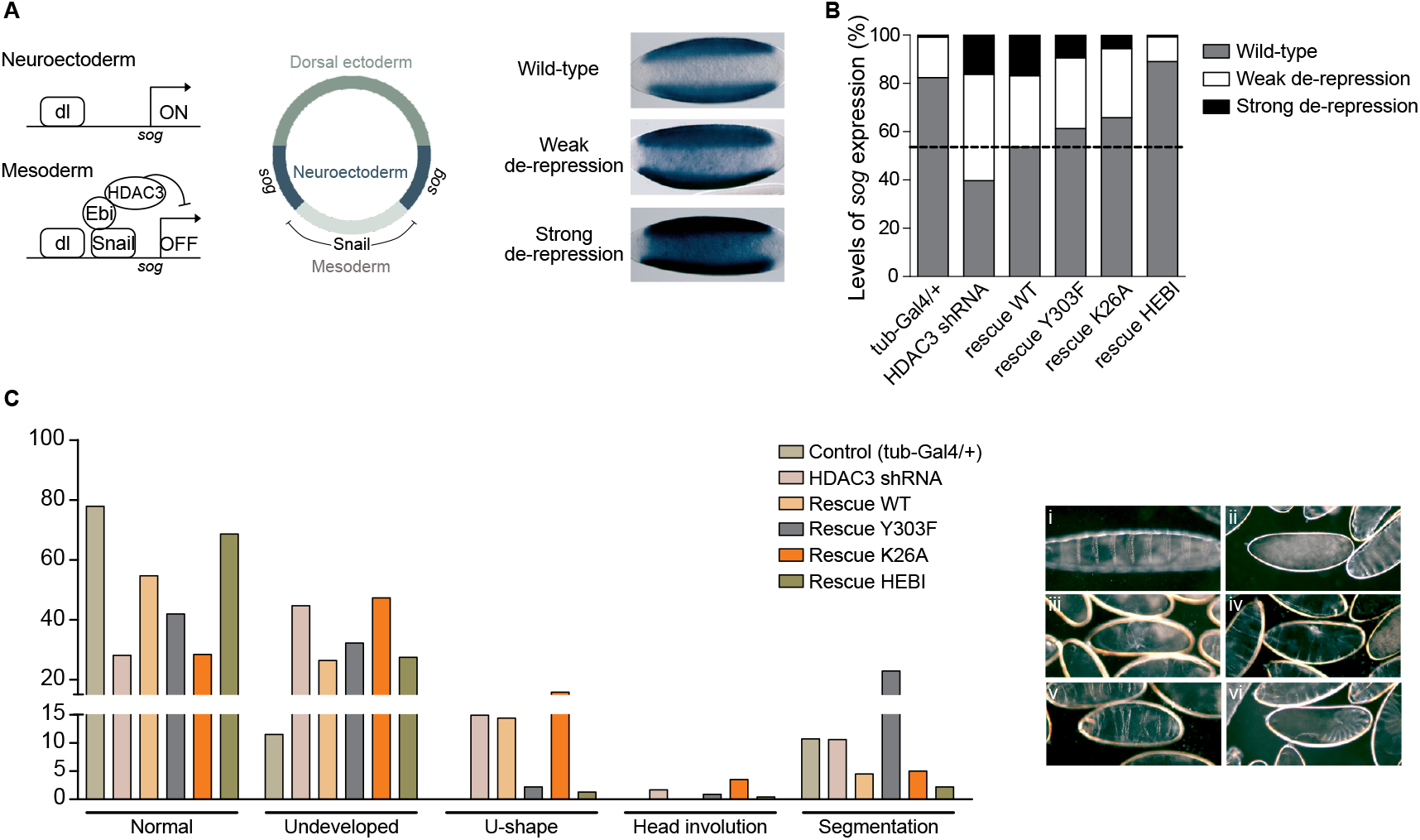
Unique functions of HDAC3 mutants in embryonic patterning. **A)** The *sog* gene is activated by Dorsal protein (dl) in the neuroectoderm of early embryos and repressed by Snail with the help of Ebi/HDAC3 in the mesoderm. In situ hybridization shows the Wild-type expression pattern of *sog* restricted to the the neuroectoderm, as well as Weak and Strong de-repression in the mesoderm. Ventral views of 2-4h embryos with anterior to the left. **B)** Graph shows the percentage of embryos with Wild-type, Weak and Strong de-repression of *sog* expression in Control (tub-GAL4/+), HDAC3 shRNA and shRNA-resistant rescue genotypes (n=199-254). **C)** Cuticle phenotypes in the depicted genotypes (n=122-317). Pictures with representative examples of embryos showing a normal cuticle phenotype (i), undeveloped (ii, iii), U-shaped (iv), segmentation (v) and head involution (vi) defects.

To further investigate the embryo phenotype we performed cuticle preparations on embryos from the different genotypes. Forty-five % of HDAC3 shRNA-depleted embryos did not develop cuticle, 28% showed a normal cuticle phenotype, 15% were U-shaped, 10% had segmentation defects, and head involution was defective in less than 2% (Fig. 3C). Consistent with the hatching rate phenotype, wild-type rescued embryos increased the fraction with a normal cuticle from 28% to 55%. The K26A rescued embryos had very similar phenotypes to the HDAC3 shRNA knock-down embryos, whereas 42% normal cuticle in Y303F rescued embryos was intermediate to knock-down and wild-type rescue (Fig. 3C). Interestingly, the frequency of segmentation defects increased to 23% in Y303F embryos, despite the presence of some wild-type HDAC3 in these embryos. This indicates that the Y303F mutation in addition to disrupting catalytic function also interferes with an additional HDAC3 activity that is needed for segmentation, since no other HDAC3 mutant showed this strong segmentation phenotype. In HEBI rescued embryos, 69% had a normal cuticle, more than in wild-type rescued embryos (Fig. 3C). Whereas embryos that did not develop cuticle were similar between HEBI and wild-type rescued, the number of embryos with a U-shaped phenotype were reduced from 14% in wild-type to 1% in HEBI-rescued embryos (Fig. 3C). The cuticle phenotype analysis shows that the different HDAC3 substitutions generate different embryonic phenotypes, indicating that there are unique functions associated with these amino acids in HDAC3.

### Separation of repressor and activator functions in HDAC3

To gain further insight into the embryonic functions of HDAC3 we generated RNA-seq data from control and HDAC3 shRNA knock-down embryos, and from knock-down embryos with shRNA-resistant wild-type, Y303F, K26A, or HEBI rescue transgenes in four biological replicates (Fig. 4, Fig. S3). Although HDAC3 is believed to function mainly as a repressor, we found that more genes were down-regulated (n=830, fold change ≥2, FDR 0.05) than up-regulated (n=179) in knock-down embryos (Fig. 4A, B, Fig. S3A). Interestingly, down-regulated genes were less strongly expressed in control embryos than most up-regulated genes, which were already highly expressed in control embryos and further increased in HDAC3 knock-down embryos (Fig. 4A). The majority of these were rescued by the wild-type HDAC3 shRNA-resistant transgene (95% of down-regulated and 93% of up-regulated genes), demonstrating little off-target effects in shRNA knock-down embryos (Fig. 4B).

**Figure 4:**
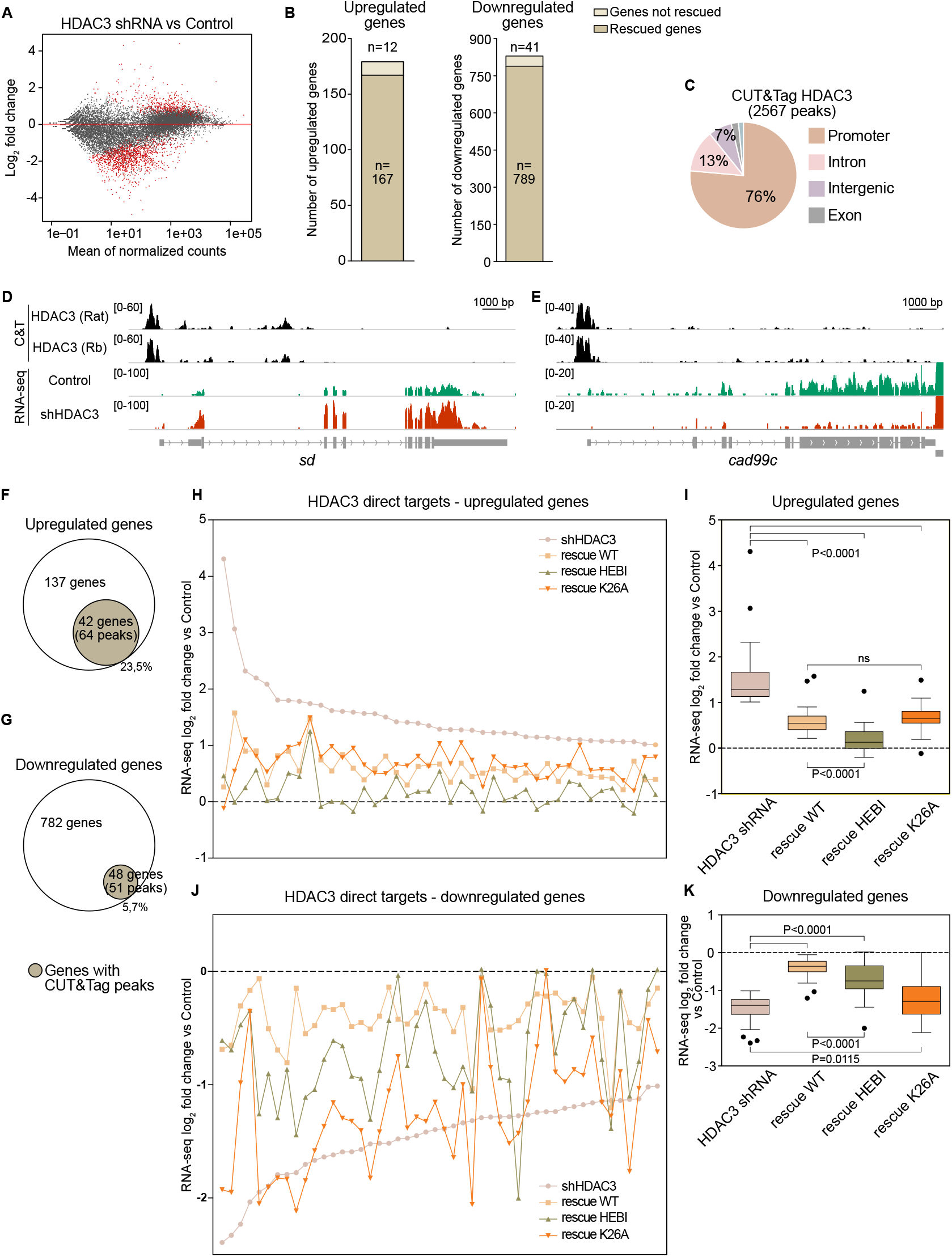
HDAC3^HEBI^ improves repressor activity and HDAC3^K26A^ impairs HDAC3 activator function in the embryo. **A)** MA plot of differentially expressed genes between HDAC3 shRNA and Control (tub-GAL4/+) identified by RNA-seq. **B)** Total number of differentially expressed genes between HDAC3 shRNA and Control (Up-regulated genes – 179, Down-regulated genes – 830, log2 fold change ≥1, FDR ≤0.05) and the number of genes rescued by the wild-type HDAC3 shRNA-resistant transgene. **C)** Genomic distribution of 2567 CUT&Tag peaks in common between two HDAC3 antibodies in 2-4h *w*^*1118*^ wild-type embryos.. **D, E)** Genome browser screenshots of one up-regulated (*sd*) and one down-regulated gene (*cad99c*) with tracks for HDAC3 CUT&Tag with two different antibodies (Rat and Rabbit) and for RNA-seq signal (HDAC3 shRNA and Control). **F, G)** Depiction of differentially expressed genes in HDAC3 shRNA embryos with HDAC3 CUT&Tag peaks. **H, J)** Expression fold change of individual HDAC3 direct targets in HDAC3 shRNA and in embryos rescued by wild-type or mutant HDAC3 relative to Control. Rescue by Y303F and an outlier in K26A is shown in Fig. S3. The dashed line intersecting the 0 on the Y axis represents the Control (Tub-Gal4/+). **I, K)** Box-plots showing the average fold expression change of HDAC3 direct target genes in HDAC3 shRNA and in embryos rescued by wild-type or mutant HDAC3 relative to Control. The dashed line intersecting the 0 on the Y axis represents the Control (Tub-Gal4/+). A two-tailed, unpaired Student’s t test was applied to compare different genotypes and error bars show standard deviation.

Since the gene expression changes can be direct or indirect, and since maternal HDAC3 shRNA knock-down also affects maternally contributed embryonic transcripts, we generated CUT&Tag data with two different HDAC3 antibodies to identify direct embryonic targets. We identified 2567 high confidence HDAC3-specific peaks in common between the two antibodies in 2-4h embryos (Fig. S3C), and most of these are located in promoters (76%) and in introns (13%) (Fig. 4C). Genome browser screenshots show examples of up-regulated and down-regulated genes (Fig. 4D, E). Only a minority of mis-expressed genes in HDAC3 knock-down embryos are bound by HDAC3 at this stage, 42 up-regulated (23%) and 48 down-regulated (6%) genes (Fig. 4F and G), suggesting that many genes are affected by maternal HDAC3 during oogenesis or are indirect targets. As a control, we performed HDAC3 CUT&Tag in wild-type and HDAC3 knock-down embryos. In this experiment, fewer wild-type peaks were detected, but they were largely depleted from knock-down embryos as only 65 out of 1142 peaks remained (Fig. S3D-F). A gene ontology analysis showed that up-regulated genes are involved in reproduction and that many direct targets are transcription factors, whereas down-regulated genes are involved in cuticle and muscle development (Table S2). Thus, HDAC3 directly regulates a few genes critical for embryonic development.

We then plotted the fold change in expression for direct targets in HDAC3 shRNA embryos and in embryos rescued by wild-type or mutant HDAC3 (Fig. 4H-K,, Fig. S3G-J, Table S1). Consistent with the hatching and cuticle results (Fig. 2 and 3), this showed that the HEBI mutant rescued most up-regulated genes to a larger extent than wild-type HDAC3 (Fig. 4H, I). This indicates that HDAC3 represses the majority of genes in the early embryo independently of its catalytic activity and independently of the interaction with SMRTER. By contrast, expression of most down-regulated genes is more efficiently rescued by wild-type than any of the mutant HDAC3s (Fig. 4J, K). Therefore, catalytic activity is required for activation but not for repression.

Interestingly, although K26A fails to rescue embryonic lethality and the cuticle phenotype, it behaves very similar to wild-type HDAC3 in rescuing the up-regulated genes (Fig. 4H, I). However, it is much poorer than wild-type at rescuing down-regulated target genes (Fig. 4J, K). Together, these results show that the HEBI mutations improve the repressor function of HDAC3, whereas K26A disrupts HDAC3 activator function.

### Targeting to the nuclear lamina or to Polycomb-repressed domains constitutes a minor form of HDAC3-dependent repression in the embryo

To investigate what mechanisms could be involved in the non-catalytic functions of HDAC3, we compared modENCODE embryonic H3K9me2 data to the gene expression changes in HDAC3 depleted embryos (Roy et al., 2010). In mammalian cells, HDAC3 has been shown to maintain some loci located within lamina-associated domains (LADs) close to the nuclear periphery (Poleshko et al., 2017). LADs are decorated with H3K9me2 (van Steensel and Belmont, 2017), and we therefore expected some of the HDAC3 targets that are enriched for this modification to be repressed by this mechanism. We found that 4 of the up-regulated genes and 6 of the down-regulated genes contain H3K9me2 in wild-type embryos (Fig. 5A). All of these up-regulated genes were rescued by both wild-type and mutant transgenes (Fig. 5B), confirming that they require a non-enzymatic HDAC3 function for their expression. A genome browser screenshot for one of them, *CG32772*, is shown in Fig. 5C. We used DNA FISH in 2-4h old embryos for this locus and it was observed to localize to the nuclear periphery in wild-type embryos (Fig. 5D). Interestingly, the locus was found at a larger distance from the periphery in HDAC3 shRNA knock-down embryos (Fig. 5D and E). By contrast, the *sog* gene locus that lacks H3K9me2 is located further from the periphery in wild-type embryos and does not shift its location towards the nuclear interior in HDAC3 knock-down embryos (Fig. S4), and is therefore repressed in an alternative catalytic-independent way. From this we conclude that nuclear lamina targeting with the help of HDAC3 is a mechanism by which around 10% of HDAC3-repressed genes could be regulated in the embryo.

**Figure 5:**
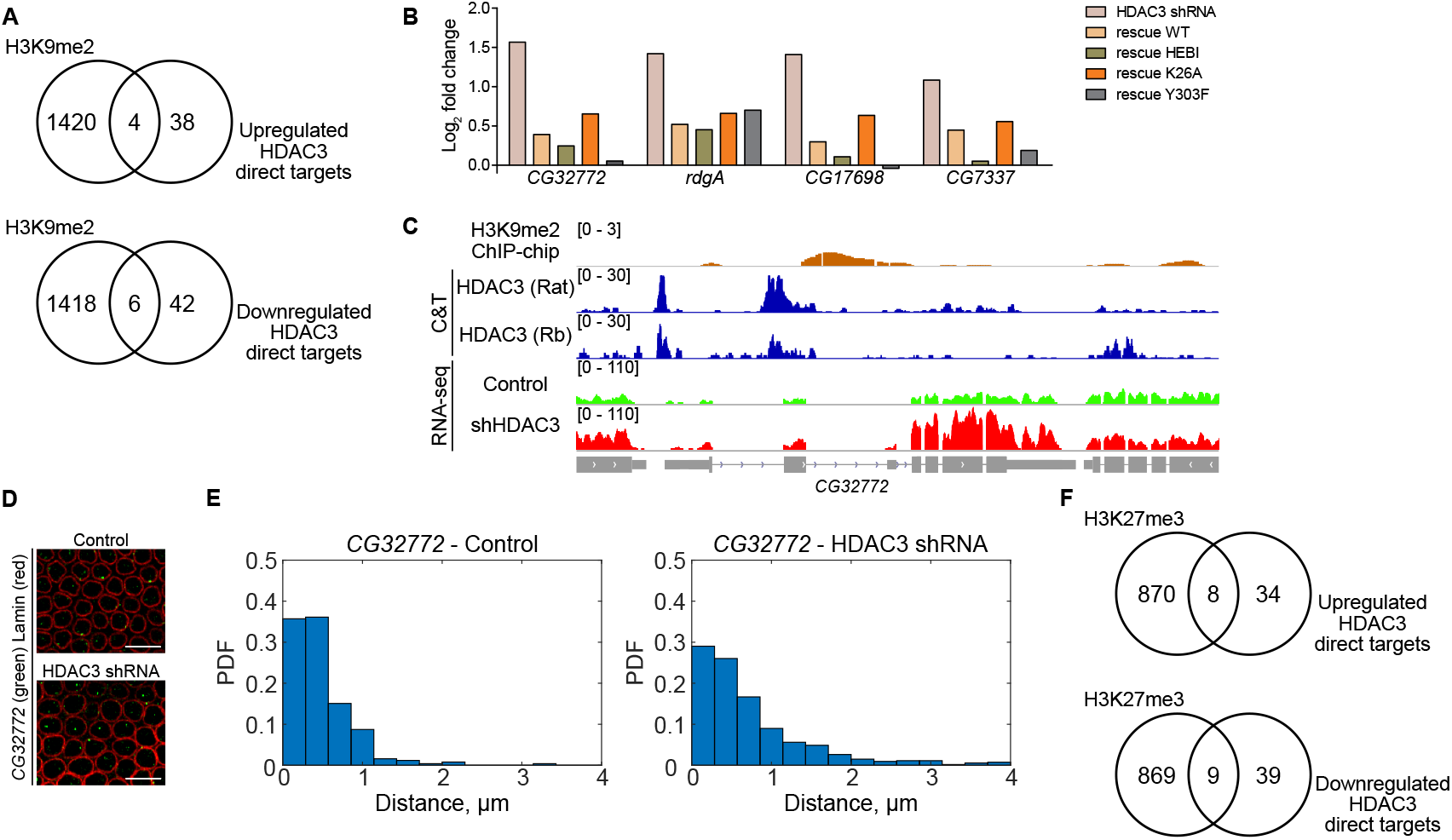
Catalytic-independent HDAC3 repression in early embryo development. **A)** Number of overlapped peaks between modENCODE embryonic H3K9me2 ChIP-chip data (Roy et al., 2010) and HDAC3 direct target genes. **B)** Expression fold change in HDAC3 shRNA and in embryos rescued by wild-type or mutant HDAC3 for up-regulated HDAC3 targets that contain H3K9me2 peaks. **C)** Genome browser screenshots of the *CG32772* locus with tracks for H3K9me2 ChIP-chip peaks, HDAC3 CUT&Tag with the two different antibodies (Rat and Rabbit) and for RNA-seq signal (HDAC3 shRNA and Control). **D)** Image of DNA FISH in 2-4h old wild-type (Control) and HDAC3 shRNA embryos using a probe for *CG32772* (green) followed by an immunostaining for Lamin (red). **E)** Measurement of the distance of the *CG32772* locus to the nuclear periphery in Control and in HDAC3 shRNA embryos, plotted as probability density functions (PDFs) where the bars in each plot sum up to one. **F)** Number of overlapped peaks between embryonic H3K27me3 ChIP-seq data (Li et al., 2014) and HDAC3 direct target genes.

Another mechanism that has been implicated in HDAC3-mediated repression is recruitment of Polycomb Group (PcG) proteins (Lewandowski et al., 2015). We therefore examined whether HDAC3-regulated genes are associated with H3K27me3 in embryos. This showed that 8 of the HDAC3-repressed genes overlap with H3K27me3 (Fig. 5F). Taken together, our data suggest that most of the HDAC3-target genes in the early embryo are repressed through an uncharacterized mode that does not require catalytic activity, targeting of loci to the nuclear lamina, or recruitment of PcG proteins.

## Discussion

The canonical function of HDAC3 is to repress target genes as part of the NCoR/SMRT complex by histone deacetylation. We find that the essential function of HDAC3 in *Drosophila* embryo development involves catalytic-dependent gene activation. A single lysine substitution, K26A, impairs catalytic activity, disrupts embryo hatching, and fails to activate target genes in the early embryo. However, it is able to repress most embryonic target genes to a similar extent as wt HDAC3. Although the mechanism by which HDAC3 activates transcription in the embryo remains unclear, studies in mammalian cells have shown that HDAC3 can activate genes in multiple contexts. In brown adipose tissue, HDAC3 associates with estrogen-related receptor alpha (ERRα) and activates the coactivator PGC-1α by deacetylation (Emmett et al., 2017). In the forebrain, HDAC3 activates some neuronal genes by its recruitment by MECP2 and deacetylation of the transcription factor FOXO3 (Nott et al., 2016). During activation of macrophages by lipopolysaccharides, HDAC3 is recruited by ATF2 and activates inflammatory genes (Chen et al., 2012; Nguyen et al., 2020). We speculate that also in the *Drosophila* embryo HDAC3 influences the activity of a transcription factor to execute gene activation.

HDAC3 has multiple deacetylase-independent functions in development and physiology. Global deletion of HDAC3 is embryonic lethal, whereas mice with mutations in both the NCoR and SMRT DAD domains that abolish HDAC3 enzymatic activity are born in expected Mendelian ratios (You et al., 2013). Furthermore, point mutants that abolish catalytic activity can partially rescue HDAC3-dependent phenotypes in the mouse liver (Sun et al., 2013). One mechanism by which HDAC3 represses transcription non-enzymatically is by recruitment of the PRC2 complex and induction of H3K27me3. In the mouse second heart field, HDAC3 silences TGFß1 expression through this mechanism (Lewandowski et al., 2015). However, we find that only around 20% of the HDAC3-repressed genes in the *Drosophila* embryo are decorated with H3K27me3.

Tethering genes to the nuclear periphery in lamina-associated domains is another non-catalytic repression mechanism. In mouse embryonic stem cells, HDAC3 tethers cardiac lineage genes to the nuclear lamina through an interaction with LAD-associated proteins such as Emerin (Demmerle et al., 2012; Poleshko et al., 2017; Somech et al., 2005; Zullo et al., 2012). We find that this is one of the mechanisms by which HDAC3 represses a subset of genes in the *Drosophila* embryo as well. However, a majority of repressed genes lack the H3K9me2 mark associated with LADs, and are therefore most likely repressed by an unknown non-enzymatic mechanism.

Interestingly, the HEBI combination of mutations improve the repressor function of HDAC3. These mutations are modelled on mammalian HDAC3 where they were shown to disrupt the interaction with NCoR/SMRT and thereby abolish catalytic activity (Sun et al., 2013). We find that corresponding mutations in *Drosophila* HDAC3 also interfere with the SMRTER interaction and impair catalytic activity. Since the HEBI mutant protein is better than wild-type at rescuing HDAC3 phenotypes, it suggests that a fraction of HDAC3 normally exists outside the SMRTER complex, is enzymatically inactive, but acts as a transcriptional repressor. Recent results in mouse macrophages found that HDAC3 associates with a subset of genes independently of NCoR/SMRT, consistent with a model of HDAC3 function outside this co-repressor complex (Nguyen et al., 2020).

Intriguingly, the enhanced repressor function of HEBI mutant HDAC3 improves survival, both when expressed from a shRNA-resistant transgene and when knocked-in at the endogenous locus. This indicates that HDAC3 has a key developmental role outside of the SMRTER complex. Interestingly, association with the NCoR/SMRT DAD-domain is allosterically regulated by the HDAC3 C-terminus, which induces a conformational change that is necessary for the interaction with the DAD-domain (Li et al., 2021a). This explains older results showing that the C-terminus is necessary for catalytic activity (Yang et al., 2002). Since the C-terminus can be phosphorylated by several kinases (Sengupta and Seto, 2004; Seo et al., 2019; Tsai and Seto, 2002; Zhang et al., 2021; Zhang et al., 2005), it suggests that the HDAC3-DAD domain interaction is regulated by signaling. Furthermore, a recent study showed that NADPH inhibits HDAC3-DAD domain complex formation by competing with IP4 for binding to HDAC3 (Li et al., 2021b). Thus, metabolic state and cell signaling regulate the balance between DAD-domain bound and unbound HDAC3, which may be a mechanism to control its transcriptional repressor activity.

## Materials & Methods

### Molecular cloning

#### HDAC3 knock in constructs

The donor construct for homologous recombination is based on the vector pP(white-STAR) from (Choi et al., 2009), in which the multiple cloning sites (MCS) were modified to insert AscI and AgeI sites at the SpeI site, and by insertion of a UAS-Rpr element for negative selection (Huang et al., 2008). The vector was named pP(whiteSTAR)Rpr13. Homologous arms of 4.4 and 3.2 kb in size were PCR amplified from genomic DNA and inserted upstream and downstream of the *white+* marker gene used for positive selection. The left arm amplicon used primers Hdac3LAfor and Hdac3LAAcc65Irev and was cloned into pGEM-T Easy, from which it was released by Acc65I digestion and cloned into the left polylinker Acc65I site in pP(whiteSTAR)Rpr13. Hdac3RAAscIfor and Hdac3RAAgeIrev primers were used to amplify the right arm, cloned into pGEM-T Easy, released with AscI and AgeI and cloned in the right polylinker of pP(whiteSTAR)Rpr13.

A 1.8 kb *HDAC3* genomic region from 263bp upstream of the HDAC3 translational start site to 66 bp downstream of the HDAC3 open reading frame was PCR amplified with primers pABCHDAC3frw and pABCHDAC3rev and cloned into the Hind III restriction site of pABC (Choi et al., 2009), which flanks the insert with attB-sites. The amino acid substitutions K26A, Y303F, or HEBI (K47E, LR160-161TK, EGAQ118-121AAAV, HNH125-127KQA) were introduced into the knock-in construct by PCR amplification of the entire vector using two phosphorylated primers, which have overhangs containing the mutations. The PCR reaction was DpnI-treated and blunt-end ligated to circularize the PCR product into a plasmid. Correct clones were identified by restriction digestion and DNA sequencing.

#### HDAC3 knock-down resistant transgenes

The *HDAC3* genomic region from 1173bp upstream of the HDAC3 TSS (transcription start site) to 771 bp downstream of the 3’ UTR was PCR amplified using primers HDAC3GR_f and HDAC3GR_r. The PCR product was cloned into the XbaI restriction site of the pattB vector (Bischof et al., 2007), resulting in the plasmid pAttB-HDAC3. A miRNA resistant sequence was introduced into pattB-HDAC3 by PCR amplification of the entire vector using two phosphorylated primers HDAC3 res_f_and HDAC3 res_r, which have overhangs containing one half of the miRNA resistant sequence each. The PCR reaction was DpnI-treated and blunt-ligated to circularize the PCR product into a plasmid. The amino acid substitutions K26A, Y303F, or HEBI (K47E, LR160-161TK, EGAQ118-121AAAV, HNH125-127KQA) were introduced into pattB-HDAC3 resistant construct by PCR amplification of the entire vector using two phosphorylated primers, which have overhangs containing the mutations. After DpnI-treatment, clones were identified by restriction digestion and DNA sequencing.

#### HA-tagged HDAC3

A HindIII restriction enzyme site was introduced into the translation termination site of the plasmid pAttB-HDAC3 by PCR amplification of the entire vector using two phosphorylated primers, HDAC3 C Hind III F and HDAC3 C Hind III R, that have overhangs containing the HindIII restriction site. Then 2 HA and 2 FLAG-tag sequences were amplified from the pRmHa-3 vector using the primers HDAC3 C tag F and HDAC3 C tag R, and cloned into the HindIII restriction site by Gibson assembly (NEB, E2611).

#### Smr constructs

The SMRTER (*Smr*) gene contains 9 exons and multiple introns of which three are long. Five pairs of primers were designed to avoid the large introns. The *Smr* genomic region was separated into 5 fragments and PCR amplified using these primers. Each fragment was cloned one by one into the XbaI restriction site of the pAC 5.1 vector that has a V5 tag, using a Gibson assembly kit (NEB, E2611). The base A was introduced into each reverse primer to form a XbaI restriction site. The final Smr full length construct deletes the large introns but contains introns between exons 3-5 and 7-9.

The Smr DAD domain is located in exons 2-3. The Smr full-lengh construct was PCR amplified using primers Smr DAD F and Smr DAD R to get the Smr DAD region. The PCR product was cloned into the XbaI restriction site of the pAC 5.1 vector.

### Fly stocks and generation of knock-out, knock-in, and miRNA resistant flies

We used a ‘ends-out’ gene targeting approach modified from (Rong and Golic, 2000) to delete the *HDAC3* gene, where the positive selection marker mini-*white*+ is flanked by attP sites and combined with negative selection using the cell death gene *reaper*. The donor construct pPWhiteSTAR Hdac3 was injected into the *w*^*1118*^ strain. Established donor lines were mapped and donor construct integrity validated as described in the supplementary materials of (Huang et al., 2008). A donor line integrated on the second chromosome was used for homologous recombination. To validate donor construct integrity the donor line was crossed to the *w-* Gal4-trap lines Gal4^477^ or Gal4^221^ that activate Rpr expression in the nervous system that leads to lethality, and to P{70FLP}10 causing mosaic eyes.

Targeting crosses were set exactly as described in (Huang et al., 2008). The 6934-hid strain contains hs-hid transgenes on the Y and balancer chromosomes, eliminating all male progeny and those female progeny carrying a hs-hid balancer chromosome after heat shock. Therefore, P{donor}/hs-FLP, hs-I-SceI females are the only genotype that survives. 50 vials each containing 30 ☿ of transgenic donor flies were mated with 30-40 ♂ of the 6934-hid strain. The crosses were maintained at room temperature and the flies flipped to fresh vials every 24 hours. Eggs in transferred vials were aged at 25°C and were heat shocked at day 2 (24-48 hours after egg-laying) and day 3 (48-72 hours after egg-laying). Heat shock was carried out at 38°C for 60-90 min in a circulating water bath.

For screening crosses 10☿ from the targeting cross were mated with *w; Gal4*^*477*^ *w-; TM2/TM6B, Tb* males to eliminate non-targeted events. Flies were flipped every couple of days 5 times and kept at 25°C. Preliminary candidates were screened by eye color based on the *white+* marker. For the mapping cross, candidates were crossed with balancer flies individually. Targeting event was confirmed by PCR with primers annealing in the *white+* gene marker and outside of the donor construct.

These knock-out flies, ΔHDAC3, where an attP-flanked mini-*white+* gene has replaced HDAC3 were used for knock-in of wild-type or mutant HDAC3 sequences. Plasmid pABC-HDAC3 containing the HDAC3 gene flanked with attB-sites was used for phiC31-mediated site-specific integration to substitute mini-*white*. Females expressing phiC31 in the germline, *y*^*1*^ *M{vas-int*.*Dm}ZH-2A w*, were crossed with ΔHDAC3 flies and the embryos injected with pABC-HDAC3. Progeny were screened for loss of mini-*white* and PCR was used to validate the integration using the primers HDAC3m knockin test F and HDAC3m knockin test R.

Bloomington stock #34778 (*y*^*1*^ *sc v*^*1*^ *sev*^*21*^; *P{TRiP*.*HMS00087}attP2*) containing a shRNA encoding a HDAC3 miRNA was used to knock-down HDAC3 in the early embryo. The *w; P{w[+mC] = matalpha4-GAL-VP16}V2H* strain (referred to as tub-Gal4) was used as maternal Gal4 driver. The pattB-HDAC3 wild-type and resistant transgenic constructs were all inserted at the attP40 landing site on the second chromosome, and combined with the shRNA shmiRNA line (#34778) on chromosome 3 and made homozygous using double balancer flies. These flies were then crossed with the tub-Gal4 driver to generate females where knock-down occurs in the female germline.

### Animal survival

The HDAC3 knock-out strain and all knock-in mutants were balanced with *TM6 Tb, Dfd-YFP*. The different mutants were selected and crossed to each other in food bottles for 2-4 days before being transferred to embryo collection cages. Embryos were collected from the cages on fruit juice agar plates supplemented with yeast and raised until 1st instar larva. The number of YFP-negative 1st instar larvae and larvae that failed to develop to 3rd instar larvae were counted.

### Embryo survival

At least 50 virgin homozygous tub-Gal4 females were placed in food bottles together with homozygous males carrying the shRNA and shRNA-resistant transgene of interest. The progeny, F1 females and males, were allowed to mate with each other in the food bottles for 2–4 days before being transferred to embryo collection cages. Embryos were collected from the cages on fruit juice agar plates supplemented with yeast. The number of embryos that could hatch was counted.

### Cuticle preparation and RNA in situ hybridization

For cuticle preparations, the embryos were aged, dechorionated in bleach, transferred to microscopic slides, cleared in lactic acid at 65°C and cuticles examined using dark-field microscopy (Wieschaus and Nüsslein-Volhard, 1988). The whole-mount embryo RNA in situ hybridization protocol was modified from (Qi et al., 2008). Probes against *sog* were generated by PCR amplification with primers where Sp6 overhangs are added, and used for in vitro transcription. We blindly counted the *sog* derepression and cuticle phenotype.

### DNA fluorescent in situ hybridization

Twelve kb gene regions centered at the TSS were selected and subdivided into 6 different fragments. DNA fragments of 1.2–1.7 kb covering 12 kb were PCR amplified from genomic DNA and labeled with Alexa Fluor 488 dye FISH Tag DNA kit (F32947, Invitrogen). DNA fluorescent in situ hybridization was performed in 2-4h old embryos based on the protocol in (Bantignies and Cavalli, 2014), and the nuclear membrane marked with anti-lamin Dm0 antibody (DSHB ADL84.12, 1:1,000) followed by a Cy5-labeled secondary antibody. For the *sog* probe, a Snail antibody was used to label the mesoderm in order to distinguish it from the neuroectoderm region.

Samples were imaged on a Nikon Ti-E microscope equipped with a sCMOS camera (Andor Technology) using a CFI Plan Apochromat Lambda 100X Oil objective (Nikon). Nuclei were segmented by first applying a median filter of size 13×13×5 pixels to suppress noise and small local variations. Then the images were segmented using thresholds calculated by Otsu’s method. Finally a morphological dilation was applied using a 5×5×3 structuring element to grow the segmentation masks slightly in order to fully cover the lamin staining. Dots were detected in AF488 and Cy5 channels using the Difference of Gaussians method. Locations were further refined by replacing each pixel coordinate by the local 3D centre of mass. For Cy5 this gave us a quite dense sampling of the lamin staining. For each point in a nucleus we reported the shortest distance to lamin (represented by any dot in Cy5) after rescaling the pixel coordinates by the pixel size. For the plots in Fig. 5 and Fig. S4 we presented the data as probability density functions (PDFs) where the bars in each plot sum up to one.

### HDAC assay

*Drosophila* S2 cells were transfected with HA-tagged HDAC3, then lysed in lysis buffer (20 mM Tris pH7.5, 150 mM NaCl, 1% NP40) containing protease inhibitors. Lysate was added to pre-washed HA-conjugated beads (Pierce anti-HA magnetic beads, catalog#88837, Thermo Scientific) and incubated at 4°C overnight. Immunoprecipitates were washed 3 times with TBST buffer and eluted with 90 ul HA peptide solution (final concentration 2mg/ml). 60 ul elution buffer was used in a HDAC assay kit (Active Motif, catalogue #56200) following the manufacturer’s instructions. Twelve ul elution buffer was separated on an 10% SDS–PAGE gel and transferred to PVDF membrane and probed with a rat anti-HDAC3 antibody diluted 1:1000 (Qi et al., 2008), which was detected with IRDye® 680RD Secondary Antibodies (LiCoR, 926-68076) using Li-CoR machine to quantify HA-tagged HDAC3. The enzyme activity was normalized to HA-tagged HDAC3 protein concentration.

### Co-immunoprecipitation

Co-immunoprecipitation was performed by preparing protein extracts from S2 cells transfected with HA-tagged HDAC3 and V5-tagged Smr full-length or DAD domain. V5 antibody (Invitrogen) was added to samples for immunoprecipitation with protein A/G magnetic beads. Protein samples were separated on an 10% SDS–PAGE gel, transferred to a PVDF membrane and probed with an HA antibody (3F10, Roche) followed by a HRP-coupled secondary antibody and enhanced chemiluminescence (ECL, GE Healthcare).

### Western blot

HDAC3 mutant larvae, obtained from 0-4h embryos and raised at 25°C for 3 days, and 2-4h knock-down embryos were harvested and directly lysed in RIPA buffer. Protein was separated by 10% SDS-PAGE gel and transferred to a PVDF membrane. The membrane was then probed with rat anti-HDAC3 (Qi et al., 2008), and anti-actin (Invitrogen, #MA5-11869) antibodies, followed by horseradish peroxidase conjugated secondary antibodies. Protein bands were visualized using enhanced chemiluminescence reagents (GE Healthcare). The protein band intensities were quantified using Image J Software using actin as a loading control.

### RNA-seq

HDAC3 knock-down and rescued 2-4h embryos were collected in biological quadruplicates and total RNA was extracted using Trizol (Invitrogen). For each biological replicate 20 ul embryos were collected. Libraries were prepared using an Illumina TruSeq kit at the Bioinformatics and Expression Analysis (BEA) facility at Karolinska Institutet and sequenced on a NextSeq 500.

### CUT&Tag

Wild-type *(w*^*1118*^) and HDAC3 shRNA embryos were collected on applejuice agar plates supplemented with yeast for two hours and aged an additional two hours. Following dechorionation, crude nuclei were isolated from around 200 embryos by homogenization using a glass douncer and a pestle in Nuclear Extraction buffer (20 mM HEPES pH 7.9, 10 mM KCl, 0.5 mM spermidine, 0,1% Triton X-100, 20% glycerol) (Hainer and Fazzio, 2019) and centrifugation at 700 g for 10 min. The obtained nuclear pellet was resuspended in 200 µL of Nuclear Extraction buffer and incubated with 20 uL of BioMag Plus Concanavalin A beads (Polysciences) for 10 min at 4 °C. The resulting nuclei/beads complex was incubated with 2 µL of primary antibody (rat or rabbit anti-HDAC3) in 200 µL of Antibody buffer (Kaya-Okur et al., 2019), overnight at 4 °C. Afterward, the experimental procedure was followed as previously described (Kaya-Okur et al., 2019), using pA-Tn5 produced at the Protein Science Facility, Karolinska Institutet and assembled with Mosaic End double-stranded (MEDS) oligonucleotide adaptors in the lab, according to (Kaya-Okur et al., 2019). The loaded pA-Tn5 was used at a 1:500 dilution. Tagmented DNA was purified with the DNA Clean & Concentrator-5 kit (D4014, Zymo Research) and amplified using PhusionHigh-Fidelity PCR Master Mix with GC Buffer (NEB) and 1.25 mM of Universal i5 primer and 1.25 mM of uniquely barcoded i7 primers for each sample (Buenrostro et al., 2015). PCR conditions were 72 °C for 5 min; 98 °C for 30 sec; thermocycling for 13 cycles at 98 °C for 10 sec and 63 °C for 10 sec; and one final step at 72 °C for 1 min. Following amplification, libraries were purified with Agencourt AMPure XP beads (Beckman Coulter) using a 1.1:1 volume of beads to sample. Libraries were sequenced (paired-end, 2×75 bp) in the NextSeq 550 Sequencing platform (Illumina) at BEA core facility, Stockholm.

### Bioinformatics

#### Differential expression analysis

RNA-seq reads were mapped to the *Drosophila melanogaster* (dm6) genome assembly using STAR and default parameters (Dobin et al., 2013). RNA-seq differential expression analysis of HDAC3 shRNA or shRNA-resistant rescue genotypes (rescue WT, rescue Y303F, rescue K26A, rescue HEBI) vs Control (tub-GAL4/+) was performed using DEseq2 (Love et al., 2014), with count tables generated from featureCounts (Liao et al., 2014). From the differentially expressed genes that were identified, we selected the ones with FDR<5% and fold change ≥ 2 (Table S?) for further analysis. Enriched GO terms among the differentially expressed genes were identified using DAVID (david.ncifcrf.gov).

#### CUT&Tag mapping and peak calling

Adaptor sequences in CUT&Tag paired-end reads were trimmed using Trim Galore! (Galaxy Tool Shed) and the trimmed reads were mapped to the *Drosophila melanogaster* (dm6) genome assembly using Bowtie2 (Langmead and Salzberg, 2012). bigWig coverage tracks were generated by normalization to the effective genome size (dm6). Peak calling of the CUT&Tag reads was performed using MACS2 (Feng et al., 2012), and overlapped peaks between different replicates/antibodies were identified using the Join tool (Galaxy Tool Shed). The obtained peaks were annotated with ChIPseeker.

## Supporting information

Table of differentially expressed genes

Table of GO terms

## Acknowledgements

We would like to thank the core facility for Bioinformatics and Expression Analysis (BEA) at Novum, which is supported by the board of research at the Karolinska Institute and the research committee at the Karolinska hospital for help with sequencing. This work was funded by grants from the Swedish Cancer Society (Cancerfonden) and the Swedish Research Council (Vetenskapsrådet) to M.M and a fellowship from the Sven and Lilly Lawski foundation to M.T.

## Author contributions

M.T. performed HDAC assay and co-IP experiments, generated and analyzed knock-down embryos, performed RNA-seq and DNA FISH, I.R. performed bioinformatic analyses, S.B. performed CUT&Tag, O.S. generated knock-in flies, L.X, E.W. and M.B. analyzed DNA FISH data, X.L and H.W performed survival assay. M.T. and M.M. designed experiments and wrote the manuscript.

## References

Achary, B.G., Campbell, K.M., Co, I.S., and Gilmour, D.S. (2014). RNAi screen in Drosophila larvae identifies histone deacetylase 3 as a positive regulator of the hsp70 heat shock gene expression during heat shock. Biochim Biophys Acta 1839, 355–363.

Bantignies, F., and Cavalli, G. (2014). Topological organization of Drosophila Hox genes using DNA fluorescent in situ hybridization. Methods Mol Biol 1196, 103–120.

Bhaskara, S., Chyla, B.J., Amann, J.M., Knutson, S.K., Cortez, D., Sun, Z.W., and Hiebert, S.W. (2008). Deletion of histone deacetylase 3 reveals critical roles in S phase progression and DNA damage control. Mol Cell 30, 61–72.

Bischof, J., Maeda, R.K., Hediger, M., Karch, F., and Basler, K. (2007). An optimized transgenesis system for Drosophila using germ-line-specific phiC31 integrases. Proc Natl Acad Sci U S A 104, 3312–3317.

Buenrostro, J.D., Wu, B., Chang, H.Y., and Greenleaf, W.J. (2015). ATAC-seq: A Method for Assaying Chromatin Accessibility Genome-Wide. Curr Protoc Mol Biol 109, 21 29 21-21 29 29.

Chen, X., Barozzi, I., Termanini, A., Prosperini, E., Recchiuti, A., Dalli, J., Mietton, F., Matteoli, G., Hiebert, S., and Natoli, G. (2012). Requirement for the histone deacetylase Hdac3 for the inflammatory gene expression program in macrophages. Proc Natl Acad Sci U S A 109, E2865–2874.

Choi, C.M., Vilain, S., Langen, M., Van Kelst, S., De Geest, N., Yan, J., Verstreken, P., and Hassan, B.A. (2009). Conditional mutagenesis in Drosophila. Science 324, 54.

Codina, A., Love, J.D., Li, Y., Lazar, M.A., Neuhaus, D., and Schwabe, J.W. (2005). Structural insights into the interaction and activation of histone deacetylase 3 by nuclear receptor corepressors. Proc Natl Acad Sci U S A 102, 6009–6014.

Cramer, P. (2019). Organization and regulation of gene transcription. Nature 573, 45–54.

Demmerle, J., Koch, A.J., and Holaska, J.M. (2012). The nuclear envelope protein emerin binds directly to histone deacetylase 3 (HDAC3) and activates HDAC3 activity. J Biol Chem 287, 22080–22088.

Emmett, M.J., and Lazar, M.A. (2019). Integrative regulation of physiology by histone deacetylase 3. Nat Rev Mol Cell Biol 20, 102–115.

Emmett, M.J., Lim, H.W., Jager, J., Richter, H.J., Adlanmerini, M., Peed, L.C., Briggs, E.R., Steger, D.J., Ma, T., Sims, C.A., et al. (2017). Histone deacetylase 3 prepares brown adipose tissue for acute thermogenic challenge. Nature 546, 544–548.

Feng, J., Liu, T., Qin, B., Zhang, Y., and Liu, X.S. (2012). Identifying ChIP-seq enrichment using MACS. Nat Protoc 7, 1728–1740.

Guenther, M.G., Barak, O., and Lazar, M.A. (2001). The SMRT and N-CoR corepressors are activating cofactors for histone deacetylase 3. Mol Cell Biol 21, 6091–6101.

Huang, J., Zhou, W., Watson, A.M., Jan, Y.N., and Hong, Y. (2008). Efficient ends-out gene targeting in Drosophila. Genetics 180, 703–707.

Kaya-Okur, H.S., Wu, S.J., Codomo, C.A., Pledger, E.S., Bryson, T.D., Henikoff, J.G., Ahmad, K., and Henikoff, S. (2019). CUT&Tag for efficient epigenomic profiling of small samples and single cells. Nat Commun 10, 1930.

Kim, Y.H., Marhon, S.A., Zhang, Y., Steger, D.J., Won, K.J., and Lazar, M.A. (2018). Rev-erbalpha dynamically modulates chromatin looping to control circadian gene transcription. Science 359, 1274–1277.

Knutson, S.K., Chyla, B.J., Amann, J.M., Bhaskara, S., Huppert, S.S., and Hiebert, S.W. (2008). Liver-specific deletion of histone deacetylase 3 disrupts metabolic transcriptional networks. EMBO J 27, 1017–1028.

Koe, C.T., Li, S., Rossi, F., Wong, J.J., Wang, Y., Zhang, Z., Chen, K., Aw, S.S., Richardson, H.E., Robson, P., et al. (2014). The Brm-HDAC3-Erm repressor complex suppresses dedifferentiation in Drosophila type II neuroblast lineages. Elife 3, e01906.

Langmead, B., and Salzberg, S.L. (2012). Fast gapped-read alignment with Bowtie 2. Nat Methods 9, 357–359.

Lewandowski, S.L., Janardhan, H.P., and Trivedi, C.M. (2015). Histone Deacetylase 3 Coordinates Deacetylase-independent Epigenetic Silencing of Transforming Growth Factor-beta1 (TGF-beta1) to Orchestrate Second Heart Field Development. J Biol Chem 290, 27067–27089.

Li, J., Guo, C., Rood, C., and Zhang, J. (2021a). A C terminus-dependent conformational change is required for HDAC3 activation by nuclear receptor corepressors. J Biol Chem 297, 101192.

Li, W., Kou, J., Qin, J., Li, L., Zhang, Z., Pan, Y., Xue, Y., and Du, W. (2021b). NADPH levels affect cellular epigenetic state by inhibiting HDAC3-Ncor complex. Nat Metab 3, 75–89.

Li, X.Y., Harrison, M.M., Villalta, J.E., Kaplan, T., and Eisen, M.B. (2014). Establishment of regions of genomic activity during the Drosophila maternal to zygotic transition. Elife 3, e03737.

Liao, Y., Smyth, G.K., and Shi, W. (2014). featureCounts: an efficient general purpose program for assigning sequence reads to genomic features. Bioinformatics 30, 923–930.

Lombardi, P.M., Cole, K.E., Dowling, D.P., and Christianson, D.W. (2011). Structure, mechanism, and inhibition of histone deacetylases and related metalloenzymes. Curr Opin Struct Biol 21, 735–743.

Lonard, D.M., and O’Malley B W. (2007). Nuclear receptor coregulators: judges, juries, and executioners of cellular regulation. Mol Cell 27, 691–700.

Love, M.I., Huber, W., and Anders, S. (2014). Moderated estimation of fold change and dispersion for RNA-seq data with DESeq2. Genome Biol 15, 550.

Lv, W.W., Wei, H.M., Wang, D.L., Ni, J.Q., and Sun, F.L. (2012). Depletion of histone deacetylase 3 antagonizes PI3K-mediated overgrowth of Drosophila organs through the acetylation of histone H4 at lysine 16. J Cell Sci 125, 5369–5378.

McHugh, C.A., Chen, C.K., Chow, A., Surka, C.F., Tran, C., McDonel, P., Pandya-Jones, A., Blanco, M., Burghard, C., Moradian, A., et al. (2015). The Xist lncRNA interacts directly with SHARP to silence transcription through HDAC3. Nature 521, 232–236.

Millard, C.J., Watson, P.J., Celardo, I., Gordiyenko, Y., Cowley, S.M., Robinson, C.V., Fairall, L., and Schwabe, J.W. (2013). Class I HDACs share a common mechanism of regulation by inositol phosphates. Mol Cell 51, 57–67.

Nguyen, H.C.B., Adlanmerini, M., Hauck, A.K., and Lazar, M.A. (2020). Dichotomous engagement of HDAC3 activity governs inflammatory responses. Nature 584, 286–290.

Nott, A., Cheng, J., Gao, F., Lin, Y.T., Gjoneska, E., Ko, T., Minhas, P., Zamudio, A.V., Meng, J., Zhang, F., et al. (2016). Histone deacetylase 3 associates with MeCP2 to regulate FOXO and social behavior. Nat Neurosci 19, 1497–1505.

Poleshko, A., Shah, P.P., Gupta, M., Babu, A., Morley, M.P., Manderfield, L.J., Ifkovits, J.L., Calderon, D., Aghajanian, H., Sierra-Pagan, J.E., et al. (2017). Genome-Nuclear Lamina Interactions Regulate Cardiac Stem Cell Lineage Restriction. Cell 171, 573–587 e514.

Qi, D., Bergman, M., Aihara, H., Nibu, Y., and Mannervik, M. (2008). Drosophila Ebi mediates Snail-dependent transcriptional repression through HDAC3-induced histone deacetylation. EMBO J 27, 898–909.

Rong, Y.S., and Golic, K.G. (2000). Gene targeting by homologous recombination in Drosophila. Science 288, 2013–2018.

Roy, S., Ernst, J., Kharchenko, P.V., Kheradpour, P., Negre, N., Eaton, M.L., Landolin, J.M., Bristow, C.A., Ma, L., Lin, M.F., et al. (2010). Identification of functional elements and regulatory circuits by Drosophila modENCODE. Science 330, 1787–1797.

Sengupta, N., and Seto, E. (2004). Regulation of histone deacetylase activities. J Cell Biochem 93, 57–67.

Seo, J., Guk, G., Park, S.H., Jeong, M.H., Jeong, J.H., Yoon, H.G., and Choi, K.C. (2019). Tyrosine phosphorylation of HDAC3 by Src kinase mediates proliferation of HER2-positive breast cancer cells. J Cell Physiol 234, 6428–6436.

Seto, E., and Yoshida, M. (2014). Erasers of histone acetylation: the histone deacetylase enzymes. Cold Spring Harb Perspect Biol 6, a018713.

Somech, R., Shaklai, S., Geller, O., Amariglio, N., Simon, A.J., Rechavi, G., and Gal-Yam, E.N. (2005). The nuclear-envelope protein and transcriptional repressor LAP2beta interacts with HDAC3 at the nuclear periphery, and induces histone H4 deacetylation. J Cell Sci 118, 4017–4025.

Sun, Z., Feng, D., Fang, B., Mullican, S.E., You, S.H., Lim, H.W., Everett, L.J., Nabel, C.S., Li, Y., Selvakumaran, V., et al. (2013). Deacetylase-independent function of HDAC3 in transcription and metabolism requires nuclear receptor corepressor. Mol Cell 52, 769–782.

Szigety, K.M., Liu, F., Yuan, C.Y., Moran, D.J., Horrell, J., Gochnauer, H.R., Cohen, R.N., Katz, J.P., Kaestner, K.H., Seykora, J.T., et al. (2020). HDAC3 ensures stepwise epidermal stratification via NCoR/SMRT-reliant mechanisms independent of its histone deacetylase activity. Genes Dev 34, 973–988.

Tsai, S.C., and Seto, E. (2002). Regulation of histone deacetylase 2 by protein kinase CK2. J Biol Chem 277, 31826–31833.

van Steensel, B., and Belmont, A.S. (2017). Lamina-Associated Domains: Links with Chromosome Architecture, Heterochromatin, and Gene Repression. Cell 169, 780–791.

Verdin, E., and Ott, M. (2015). 50 years of protein acetylation: from gene regulation to epigenetics, metabolism and beyond. Nat Rev Mol Cell Biol 16, 258–264.

Watson, P.J., Fairall, L., Santos, G.M., and Schwabe, J.W. (2012). Structure of HDAC3 bound to co-repressor and inositol tetraphosphate. Nature 481, 335–340.

Wieschaus, E., and Nüsslein-Volhard, C. (1988). Looking at embryos. In Drosophila: A Practical Approach, D.B. Roberts, ed. (Oxford University Press), pp. 199–227.

Yang, W.M., Tsai, S.C., Wen, Y.D., Fejer, G., and Seto, E. (2002). Functional domains of histone deacetylase-3. J Biol Chem 277, 9447–9454.

Yin, H., Kang, Z., Zhang, Y., Gong, Y., Liu, M., Xue, Y., He, W., Wang, Y., Zhang, S., Xu, Q., et al. (2021). HDAC3 controls male fertility through enzyme-independent transcriptional regulation at the meiotic exit of spermatogenesis. Nucleic Acids Res 49, 5106–5123.

You, S.H., Lim, H.W., Sun, Z., Broache, M., Won, K.J., and Lazar, M.A. (2013). Nuclear receptor co-repressors are required for the histone-deacetylase activity of HDAC3 in vivo. Nat Struct Mol Biol 20, 182–187.

Zhang, F., Qi, L., Feng, Q., Zhang, B., Li, X., Liu, C., Li, W., Liu, Q., Yang, D., Yin, Y., et al. (2021). HIPK2 phosphorylates HDAC3 for NF-kappaB acetylation to ameliorate colitis-associated colorectal carcinoma and sepsis. Proc Natl Acad Sci U S A 118.

Zhang, L., He, X., Liu, L., Jiang, M., Zhao, C., Wang, H., He, D., Zheng, T., Zhou, X., Hassan, A., et al. (2016). Hdac3 Interaction with p300 Histone Acetyltransferase Regulates the Oligodendrocyte and Astrocyte Lineage Fate Switch. Dev Cell 36, 316–330.

Zhang, X., Ozawa, Y., Lee, H., Wen, Y.D., Tan, T.H., Wadzinski, B.E., and Seto, E. (2005). Histone deacetylase 3 (HDAC3) activity is regulated by interaction with protein serine/threonine phosphatase 4. Genes Dev 19, 827–839.

Zhu, C.C., Bornemann, D.J., Zhitomirsky, D., Miller, E.L., O’Connor, M.B., and Simon, J.A. (2008). Drosophila histone deacetylase-3 controls imaginal disc size through suppression of apoptosis. PLoS Genet 4, e1000009.

Zullo, J.M., Demarco, I.A., Pique-Regi, R., Gaffney, D.J., Epstein, C.B., Spooner, C.J., Luperchio, T.R., Bernstein, B.E., Pritchard, J.K., Reddy, K.L., et al. (2012). DNA sequence-dependent compartmentalization and silencing of chromatin at the nuclear lamina. Cell 149, 1474–1487.

