## Supplemental Figures for "Separation of transcriptional repressor and activator functions in HDAC3"

Figure S1

**A K26A mutant**

```
                                                     K26A
HDAC3_Dmel     -----MTDRRVSYFYNADVGNFHYGAGHPMKPQRLAVTHSLVMNYGLHKKMKIYRPYKAS   55
HDAC3_Human    ------MAKTVAYFYDPDVGNFHYGAGHPMKPHRLALTHSLVLHYGLYKKMIVFKPYQAS   54
HDAC1_Human    MAQTQGTRRKVCYYYDGDVGNYYYGQGHPMKPHRIRMTHNLLLNYGLYRKMEIYRPHKAN   60
HDAC1_Dmel     --MQSHSKKRVCYYYDSDIGNYYYGQGHPMKPHRIRMTHNLLLNYGLYRKMEIYRPHKAT   58
```

**B Y303F mutant**

```
                     Y303F
HDAC3_Dmel     LVVGGGGYTLRNVARCWTHETSLLVDQDIENDLPATEYYDFFAPDFTLHPEINSRQDNAN   355
HDAC3_Human    LVLGGGGYTVRNVARCWTYETSLLVEEAISEELPYSEYFEYFAPDFTLHPDVSTRIENQN   350
HDAC1_Human    LMLGGGGYTIRNVARCWTYETAVALDTEIPNELPYNDYFEYFGPDFKLHISP-SNMTNQN   354
HDAC1_Dmel     LMVGGGGYTIRNVSRCWTYETSVALAVEIANELPYNDYFEYFGPDFKLHISP-SNMTNQN   352
```

**C HEBI mutant**

```
                                                                K47E
HDAC3_Dmel     -----MTDRRVSYFYNADVGNFHYGAGHPMKPQRLAVTHSLVMNYGLHKKMKIYRPYKAS   55
HDAC3_Human    ------MAKTVAYFYDPDVGNFHYGAGHPMKPHRLALTHSLVLHYGLYKKMIVFKPYQAS   54
HDAC1_Human    MAQTQGTRRKVCYYYDGDVGNYYYGQGHPMKPHRIRMTHNLLLNYGLYRKMEIYRPHKAN   60
HDAC1_Dmel     --MQSHSKKRVCYYYDSDIGNYYYGQGHPMKPHRIRMTHNLLLNYGLYRKMEIYRPHKAT   58

                   LR60TK
HDAC3_Dmel     AQDMLRFHSDEYIAYLQQVTPQNIQCNSVAYTKYLAHFSVGEDCPVFDGLFDFCAMYTGA   115
HDAC3_Human    QHDMCRFHSEDYIDFLQRVSPTNMQ----GFTKSLNAFNVGDDCPVFPGLFEFCSRYTGA   110
HDAC1_Human    AEEMTKYHSDDYIKFLRSIRPDNMS----EYSKQMQRFNVGEDCPVFDGLFEFCQLSTGG   116
HDAC1_Dmel     ADEMTKFHSDEYVRFLRSIRPDNMS----EYNKQMQRFNVGEDCPVFDGLYEFCQLSAGG   114

          EGAQ118AAAV  HNH125KQA
HDAC3_Dmel     SLEGAQKLNHNHSDICINWSGGLHHAKKFEASGFCYVNDIVIGILELLKYHPRVLYIDID   175
HDAC3_Human    SLQGATQLNNKICDIAINWAGGLHHAKKFEASGFCYVNDIVIGILELLKYHPRVLYIDID   170
HDAC1_Human    SVASAVKLNKQQTDIAVNWAGGLHHAKKSEASGFCYVNDIVLAILELLKYHQRVLYIDID   176
HDAC1_Dmel     SVAAAVKLNKQASEICINWGGGLHHAKKSEASGFCYVNDIVLGILELLKYHQRVLYIDID   174
```

**D**

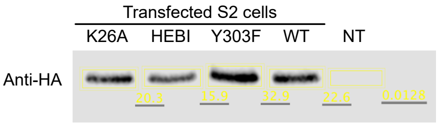

**Figure S1: Sequence alignment of *Drosophila melanogaster* and human HDAC3 and HDAC1 proteins. A-C)** *Drosophila melanogaster* HDAC3 mutations expected to interfere with catalytic activity were modelled on human HDAC3. Red rectangles indicate the amino acid substitutions in each mutant. **A)** K26A **B)** Y303F **C)** HEBI. **D)** Western-blot shows protein extracts from S2 cells transfected wild-type and mutant HA-tagged HDAC3 after immunoprecipitation with an anti-HA antibody.

Figure S2

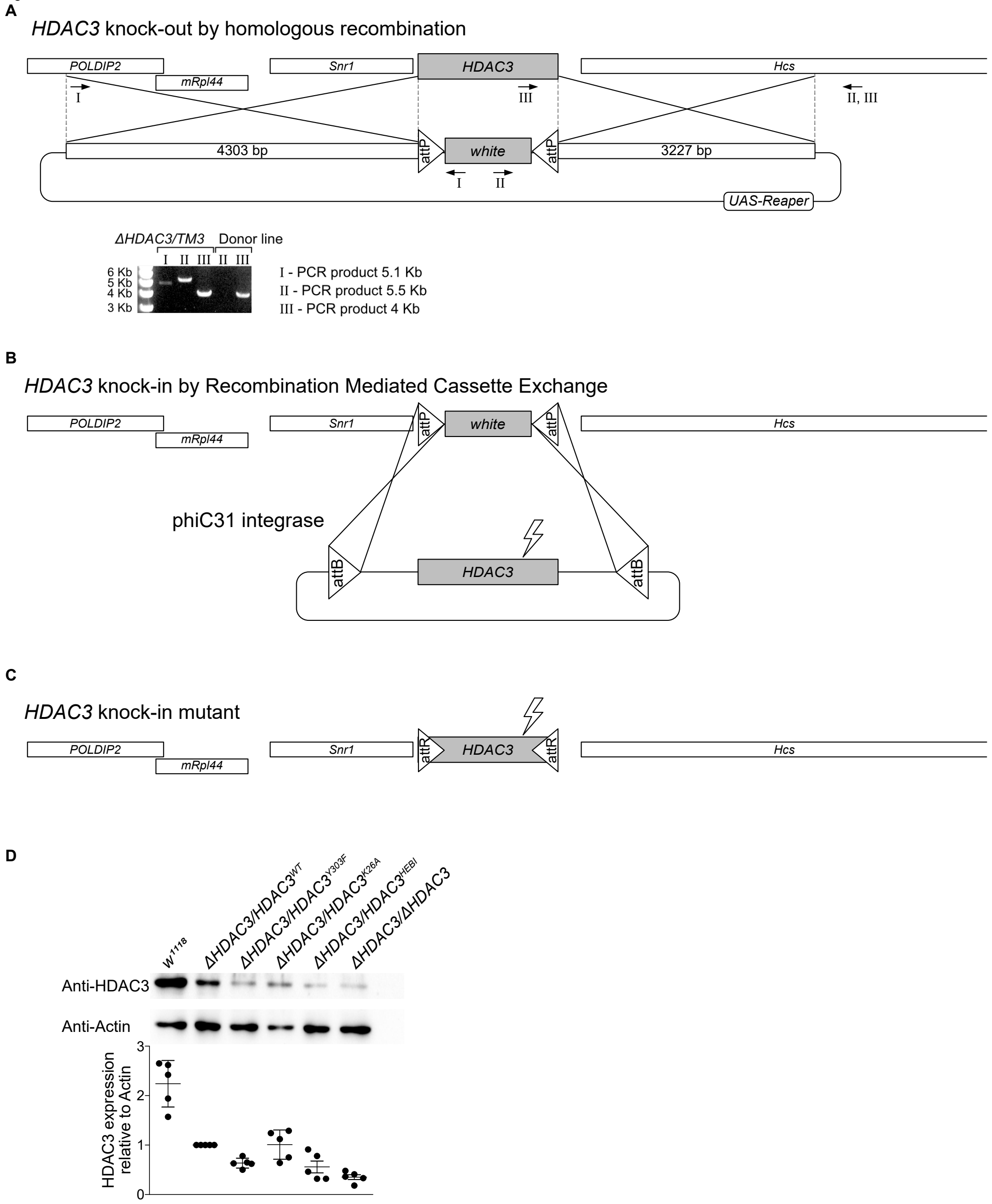

**Figure S2: Generation of HDAC3 knock-in mutant flies. A)** The HDAC3 knock-out was created by replacement of the HDAC3 transcription unit with a *mini-white* (*white*) gene flanked by attP sites by homologous recombination. I to III indicate the locations of PCR primer pairs used to confirm the recombination in the balanced ΔHDAC3 line (ΔHDAC3/TM3) and compared to the donor line before recombination. **B, C)** The HDAC3 wild-type or mutant sequences were re-introduced into the endogenous HDAC3 locus using a recombination-mediated cassette exchange

(RMCE)-based strategy with the phiC31 integrase. **D)** Western-blot from second instar larvae showing the levels of HDAC3 relative to Actin in wild-type (*$w^{1118}$*) and trans-heterozygous HDAC3 mutant animals.

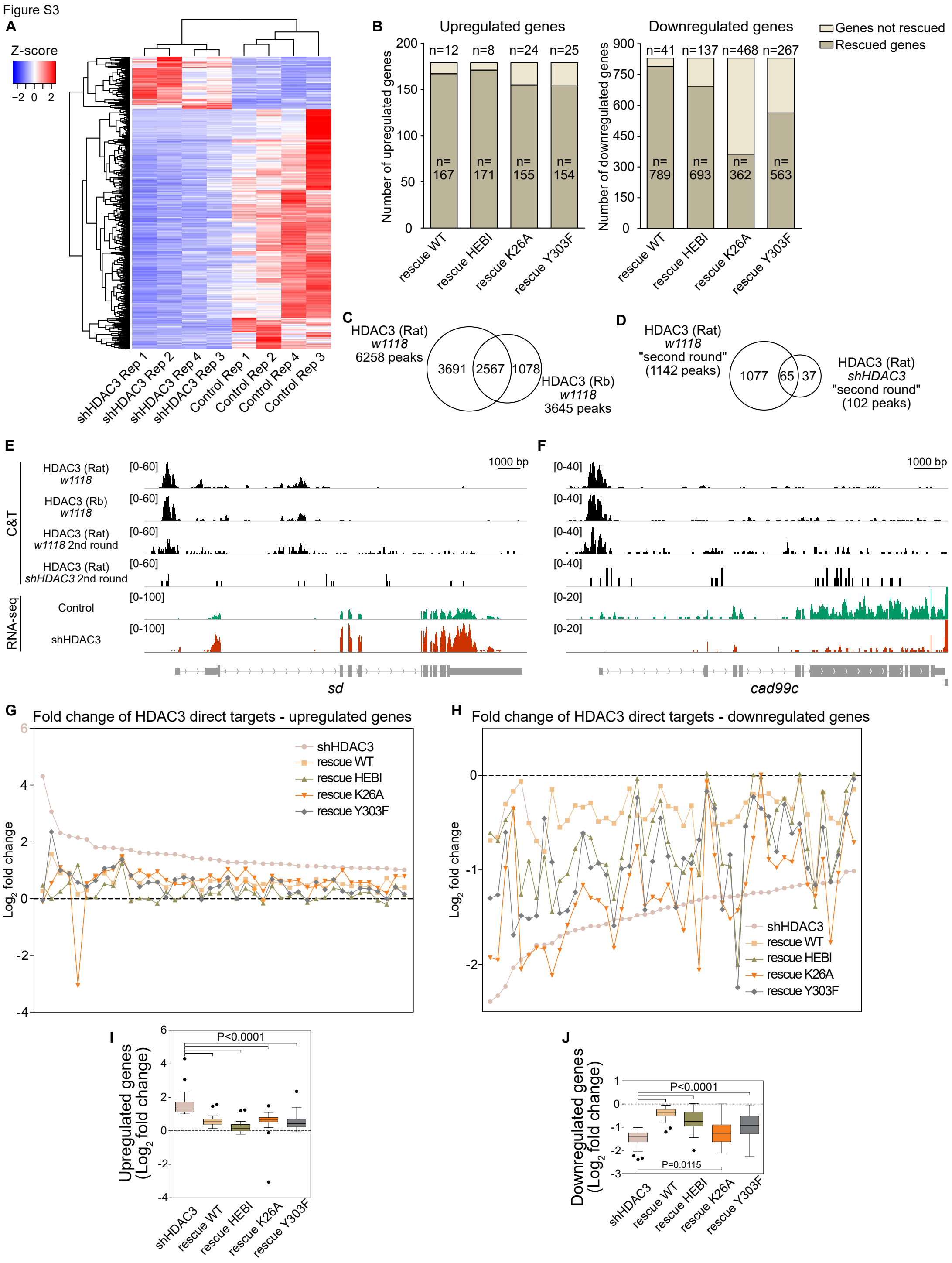

**Figure S3: RNA-seq and CUT&Tag in control and HDAC3 shRNA knock-down embryos. A)** Heatmap showing normalized expression in HDAC3 shRNA and Control replicates standardized to Z-score by subtracting the mean and dividing by the standard deviation of the significantly changed genes ($\log_2$ fold change $\geq 1$; FDR $<0.05$). **B)** Number of differentially expressed genes between HDAC3 shRNA and

Control ($\log_2$ fold change ≥1, FDR ≤0.05) and the number of genes rescued by the wild-type or mutant HDAC3 shRNA-resistant rescue transgenes. **C)** Number of total and overlapped peaks between HDAC3 CUT&Tag data generated with two different antibodies (Rat and Rabbit) in 2-4h old *w*$^{1118}$ embryos. **D)** Number of total and overlapped peaks between HDAC3 CUT&Tag in 2-4h old *w*$^{1118}$ and HDAC3 shRNA embryos. **E, F)** Genome browser screenshots of one up-regulated (*sd*) and one down-regulated gene (*cad99c*) with tracks for HDAC3 CUT&Tag with the two different antibodies (Rat and Rabbit) in 2-4h old *w*$^{1118}$ embryos, the second CUT&Tag experiment in 2-4h old *w*$^{1118}$ and HDAC3 shRNA embryos and for RNA-seq signal (HDAC3 shRNA and Control). **G, H)** Expression fold change of individual HDAC3 direct targets in HDAC3 shRNA and in embryos rescued by wild-type or mutant HDAC3 relative to Control. The dashed line intersecting the 0 on the Y axis represents the Control (Tub-Gal4). **I, J)** Box-plots showing the average fold expression change of HDAC3 direct target genes in HDAC3 shRNA and in embryos rescued by wild-type or mutant HDAC3 relative to Control.. The dashed line intersecting the 0 on the Y axis represents the Control (Tub-Gal4/+). A two-tailed, unpaired Student's t test was applied to compare different genotypes and error bars show standard deviation.

Figure S4

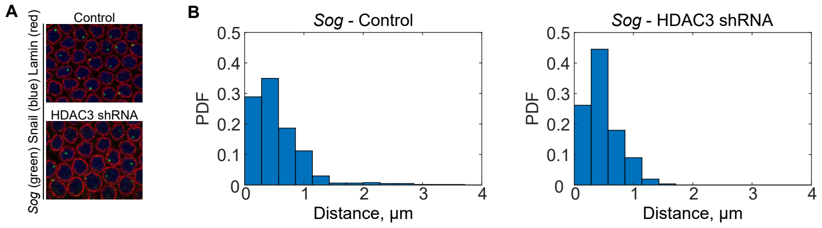

**Figure S4: The *sog* gene is not repressed by nuclear lamina targeting. A)** Image of DNA FISH in 2-4h old wild-type (Control) and HDAC3 shRNA embryos using a probe *sog* (green) followed by an immunostaining for Lamin (red) and Snail (blue). Snail staining indicates nuclei in the mesoderm. **B)** Measurement of the distance of the *sog* locus to the nuclear periphery in Control and in HDAC3 shRNA embryos.

| Primer name | Primer sequence (5' - 3') | Notes |
|---|---|---|
| **HDAC3 knock in constructs** | | |
| Hdac3LAfor | TCCCCAACGGCTGCTACC | |
| Hdac3LAAcc65Irev | GGTACCAGCGTAAGCAGTAAAGAGTATCA | |
| Hdac3RAAscIfor | GGCGCGCCTAGTGTACTAAAAATGGA | |
| Hdac3RAAgeIrev | TACCGGTGGTTTGCGAGAGTGCGAC | |
| pABCHDAC3frw | GAGAAAGCTTCGCCGCTTTAAGTTGTCTTC | |
| pABCHDAC3rev | GAGAAAGCTTACAAAATAAGCTGATACTCTTTACTGC | |
| HDAC3m knockin test F | GGACAAGCTGGTTGATGCTA | |
| HDAC3m knockin test R | AGGCCCCTCTATCAACGATT | |
| **HDAC3 knock down resistant transgenes** | | |
| HDAC3GR F | acacta**TCTAGA**GGGCCGCTGTAGTGGCTCTATG | |
| HDAC3GR R | atacaA**TCTAG**AATGCGGCGCAGAAGGAGTG | |
| HDAC3res F | TACTGTAG AATGTAGAATAAGGATTCGA | |
| HDAC3res R | TTAGCTCTCC GCCGAATCGGGCTTGTCTT | |
| HDAC3 Y303F F | TTCACCTTACGGAACGTGGC | Y303F TAC>TTC |
| HDAC3 Y303F R | GCCACCGCCTCCGACC | |
| HDAC3 K26A F | GCGCCGCAACGCCTGGCG | |
| HDAC3 K26A R | CATGGGATGCCCGGCTCCATAG | |
| HDAC3 K47E F | gaGATCTACAGGCCGTACAAGTGAG | K47E AAG>GAG |
| HDAC3 K47E R | CATTTTCTTGTGCAGTCCGTAG | |
| HDAC3 HIm F | CTGAACAAGCAAGCCAGCGACATATGCATAAACTGGTCGG | HNH125-127KQA |
| HDAC3 HIm R | CTTCACGGCGGCGGCCAGCGAGGCACCCGTGTAC | EGAQ118-121AAAV |
| HDAC3 Em F | AAGTTCCACAGCGACGAGTACATC | R61K CGC > AAG |
| HDAC3 Em R | ggtCATGTCCTGCGCACTGG | L60T CTC > ACC |
| **HA-tagged HDAC3** | | |
| HDAC3 C Hind III F | AAGCTTACTTTCTGCCGAATCGGGCTT | |
| HDAC3 C Hind III R | TAGTATTGGATAATGTAGAATAAGGATTCGA | |
| HDAC3 C tag F | AGACAAGCCCGATTCGGCAGAAAGTCGATCCTACCCCTACGATGTT | |
| HDAC3 C tag R | CCTTATTCTACATTATCCAATACTACTTGTCATCATCGTCCTTGTAGTCGAG | |
| **Smr constructs** | | |
| Smr full length F | CCAGCACAGTGGCGGCCGCTCGAGTATGGAAGCAGCTATACGCTCCA | Exon 1 |
| smr 9R-2 | GGCTTACCTTCGAAGGGCCCTCTAGACTGTGAATGGATGCTGCATTTG | |
| smr 9F-2 | CATCAAATGCAGCATCCATTCACAGCCCCCACTGGTCTTACAG | Exon 2 |
| smr 10R-2 | GGCTTACCTTCGAAGGGCCCTCTAGACTGAGGATGATAGGCCTC | |
| smr 10F-2 | CAGCACCGAGGCCTATCATCCTCAGGTGGAGGCCATTTCGCCG | Exon 3-5 |
| smr 11R-2 | GGCTTACCTTCGAAGGGCCCTCTAGACTTTGCCATCGAGGGCGAG | |
| smr 11F-2 | GAGATTCTCGCCCTCGATGGCAAAGACAAATTGGCCAGCTGCTTTG | Exon 6 |
| smr 12R-2 | GGCTTACCTTCGAAGGGCCCTCTAGACTATTAACATGATCGGCTGGC | |
| smr 12F-2 | CATCGCCAGCCGATCATGTTAATAGCACTCCGAGTCCGCATCG | Exon 7-9 |
| Smr full length R-2 | GGCTTACCTTCGAAGGGCCCTCTAGAATCTTCGTCGCTGAGTGCAT | |
| smrDAD F | CCAGCACAGTGGCGGCCGCTCGAGTATGAATGTTGCCGCCGGATTGGGTA | |
| smrDAD R | GGCTTACCTTCGAAGGGCCCTCTAGAATCCATGATCTCCTCGAGATCCGCT | |
| **DNA FISH** | | |
| sog fish f1 | TGCCCCATGTGCGACTATTA | 1666bp |
| sog fish r1 | TAAGAAAGAGCGGTCCAGGG | |
| sog fish f2 | GGGTAGCTGATGGGAATGGT | 1225bp |
| sog fish r2 | GACCATAAAGCGGCACCCAA | |
| sog fish f3 | TCTACCTGCGATTACGGGGA | 1516bp |
| sog fish r3 | GATTGCACATCGGTCGTTCG | |
| sog fish f4 | GGGTCGACAGTCCGAGAATC | 1368bp |
| sog fish r4 | TGGTAGAGCCGGCTGAGATA | |
| sog fish f5 | AAATGTTTTGGCCTCACGGC | 1346bp |
| sog fish r5 | CACACACCGGCTACTGTTCT | |
| sog fish f6 | ACTGGTCCGCATCGAATGTT | 1297bp |
| sog fish r6 | CTACACAGCGATTTGTCGCC | |
| CG32772 fish f1 | GCGTAACATAGCGTTTGGTGCTGTC | 1214bp |
| CG32772 fish r1 | ATATGGCCACAATCAGCATGGAGGG | |
| CG32772 fish f2 | AGCATGGTGGAGTTCGCCTTTGTTA | 1666bp |
| CG32772 fish r2 | CTTTGTTGCTGACTTTTGCCCCCTC | |
| CG32772 fish f3 | AATGACAGAAATTGCGCCAAGGCAG | 1475bp |
| CG32772 fish r3 | TGTGAGCGGGGGAGTAGTAGTGAAA | |
| CG32772 fish f4 | GCACGGGAATTTGCAATCCGTTCTT | 1450bp |
| CG32772 fish r4 | TTGATCTCGATCTTGATGGCCGTGG | |
| CG32772 fish f5 | ATCGGCGAGTGGACTTCCAAAACAT | 1314bp |
| CG32772 fish r5 | TCGTTTCTCTCTCCCCCAGGACATT | |
| CG32772 fish f6 | TGCCAATGAACAGCGGATACGAACT | 1585bp |
| CG32772 fish r6 | CTCTTCAACTACGGGCTGCATCACA | |
